# Putative SET-domain methyltransferases in *Cryptosporidium parvum* and histone methylation during infection

**DOI:** 10.1101/2022.03.06.483160

**Authors:** Manasi Sawant, Sadia Benamrouz-Vanneste, Dionigia Meloni, Nausicaa Gantois, Gaёl Even, Karine Guyot, Colette Creusy, Erika Duval, René Wintjens, Jonathan Weitzman, Magali Chabe, Eric Viscogliosi, Gabriela Certad

## Abstract

*Cryptosporidium parvum* is a major cause of an intestinal pathology called cryptosporidiosis which affects humans and other vertebrates. Despite being declared as a public health problem by World Health Organization (WHO) since 2006, pathogenesis caused by this parasite remains poorly understood. More recently, *C. parvum* has been linked with oncogenesis. In particular, the mechanisms involved in the processes of gene expression regulation are completely unexplored in *Cryptosporidium*. In the current study, we took the opportunity to investigate a dynamic epigenetic modification called histone lysine methylation during the life cycle of the parasite. We successfully identified putative SET-domain containing proteins, lysine methyltransferases (KMTs), which catalyze the methylation of different lysine residues. Phylogenetic analysis classified them into distinct subfamilies namely CpSET1, CpSET2, CpSET8, CpKMTox and CpAKMT. Structural analysis further characterized CpSET1, CpSET2 and CpSET8 to be histone lysine methyltransferases (HKMTs). Their functional significance was predicted by using site-specific methyl-lysine antibodies during development of the parasite (CpSET1:H3K4; CpSET2:H3K36; CpSET8:H4K20). In particular, the SET domain of CpSET8 showcased methyltransferase activity confirming the existence of functional HKMTs in *Cryptosporidium*. Moreover, the consequence of *C. parvum* infection on the host lysine methylation events highlights the inherit potential of the parasite to exploit the host epigenetic regulation to its advantage. Thus, this study is the first one to provide insights on epigenetics mechanisms occurring throughout the parasite’s life cycle and during the interaction with its host. As *Cryptosporidium* is a protozoan that significantly affects the health of both humans and animals, a better understanding of its developmental processes within the definitive host may highlight novel infection control strategies.

**Author Summary:** *Cryptosporidium* species have a very compact genome (~9.2 Mb) unlike its apicomplexan homologs such as *Toxoplasma* (~63 Mb). Moreover, the lack of large families of transcriptional factors requires them to heavily rely on chromatin remodeling components for its gene regulation. Thus, study and identification of novel elements which contribute to chromatin dynamics could assist a better understanding of the biology of this parasite. In the current study we investigated histone lysine methylation, a dynamic epigenetic modification which regulates gene activation as well as repression. More importantly, characterizing the enzymes which bring about this regulation, provides potential new druggable targets to attack the parasite.

## 1. Introduction

*Cryptosporidium* belong to the eukaryotic phylum Apicomplexa, distinct relatives of the parasites that cause malaria and toxoplasmosis (1). *Cryptosporidium* infection, also termed as cryptosporidiosis, is responsible for self-limited diarrhea in healthy immunocompetent individuals but is capable of causing life-threatening disease among those who are immunocompromised (2). Most recently, a series of epidemiological studies has reported the diarrheal disease caused by the parasite to be one of the leading causes of early childhood morbidity and mortality in developing countries, together with rotavirus, enterotoxigenic *Escherichia coli* and *Shigella* (3,4). The environmentally resilient nature of the *Cryptosporidium* oocysts allows the parasite to withstand common water treatments such as chlorination (5), and therefore to remain a major cause of waterborne outbreaks in industrialized countries as well (6,7). Even though being identified to have a significant impact on public health, no vaccine or chemoprophylactic drugs to prevent *Cryptosporidium* infection and very few chemotherapeutic options for its treatment are currently available (8).

Majority of human infections by this protozoan are caused by *Cryptosporidium hominis* and *Cryptosporidium parvum*. Thus, new opportunities to uncover targets for drugs are attributed to the availability and *in silico* analysis of genome sequences of *C. hominis* (9) and *C. parvum* (10). These two genomes are small in size (approximately 9 Mb), adenosine- and thymidine-rich (approximately 70%) and differ only at nucleotide level by 3% −5% (9,10). In parallel, *C. parvum* transcriptomes at the oocyst, excysted sporozoites and intracellular stages have been widely investigated to understand *Cryptosporidium* life cycle development (11,12) even though the asynchronous nature of the parasite life cycle has made the transcriptome interpretations challenging. Interestingly, with the availability of the technologies to genetically manipulate the parasite (13), it became possible to analyze transcriptome at specific intracellular stages *in vitro* as well as *in vivo* (14), leading to the discovery of novel targets for therapeutic interventions.

Strikingly, the compact genome of *Cryptosporidium* compared to other apicomplexan homologs (e.g approximately 63 Mb for *T. gondii*) accompanied by the lack of large families of recognizable transcription factors typically encountered in eukaryotic organisms (15), suggest major differences in the mechanisms of apicomplexan gene regulation. Accordingly, these mechanisms may depend on epigenetic events in order to control gene expression and cellular differentiation (16).

In eukaryotes, epigenetic changes include DNA methylation (17) and histone modifications such as lysine methylation (18) and acetylation (19). Apicomplexan parasites such as *Toxoplasma* and *Cryptosporidium* are known to encode putative DNA methylation enzymes but they lack detectable DNA cytosine methylation events (20). On the other hand, novel drugs targeting the enzyme histone deacetylase (HDAC) which results in hyperacetylation of their genomes causing arrest of parasite differentiation has been reported in *Toxoplasma* (21), *Plasmodium* (22) and *Cryptosporidium* (23). Along with acetylation, histone lysine methylation, a rather more sophisticated and dynamic post-translational modification previously identified to be restricted to metazoans has been extensively studied in *Toxoplasma* and *Plasmodium* (16). Lysine methyltransferases (KMTs) known to methylate Histone 4 lysine 20 (H4K20), were identified to regulate cell cycle progression in these two apicomplexan parasites (24). In *Plasmodium*, lysine methylation markers such as Histone 3 lysine 4 (H3K4) and Histone 3 lysine 9 (H3K9) have been reported to be involved in regulatory mechanisms of switching between variant surface antigens, enabling the parasite to evade the host immune response (25,26). Histone methylation is a reversible event catalyzed by histone demethylases. Jumonji-C-terminal (JmjC) domain-containing putative histone demethylases have been identified in *T. gondii, P. falciparum, Babesia bovis*, and *Theileria annulata* (27). Databases have also identified two lysine-specific demethylases (LSD)-like proteins in *T. gondii* genome (28). However, these epigenetic modifications have not been previously investigated in *Cryptosporidium* parasites.

On the other hand, pathogens usually induce alterations of their hosts employing several strategies to target cellular processes during their complex interactions with host cells (29). In this way, they can evade the barriers imposed by checkpoint responses, and can manipulate various pathways activated for cell protection in order to increase their survival and transmission. Then, epigenetic mechanisms could play a fundamental role in the dynamic of host–parasite interactions (29). As an example, lysine methylation is emerging as a versatile and dynamic post-translational modification that contributes critically to cellular differentiation programs being a pivotal dynamic event in host-pathogen interactions (30). Interestingly, *Theileria*, which causes a lymphoproliferative disease in cow manipulates the leukocyte epigenetic mechanisms favoring parasite replication and persistence leading to the host cell transformation (30). Strikingly, epidemiological and experimental studies suggest a potential link between *Cryptosporidium* infection and digestive cancer (31). Of all apicomplexan parasites, *Cryptosporidium* and *Theileria* are able to transform their mammalian hosts. However, little is known about the significance of epigenetic variations in *Cryptosporidium* development and in the parasite interactions with its host.

In the current study, we thus aim to characterize the KMTs of *Cryptosporidium* in order to identify lysine methylation events which might be involved in regulating gene expression during the life cycle of the parasite and to evaluate host epigenetic modulators potentially involved in pathogenicity and parasite-induced transformation. In parallel, this study also highlights the potential of lysine methylation modifications to be considered as targets in the development of therapeutic strategies against *Cryptosporidium* infection.

## 2. Results

### 2.1. *In silico* identification of putative KMTs of *C. parvum* and sequence comparison of their SET domains

Queries were run to retrieve KMTs encoding genes from *C. parvum* containing SET domain consensus. As a result, ten putative KMTs with a recognisable SET domain were predicted (Table 1).

**Table 1.**
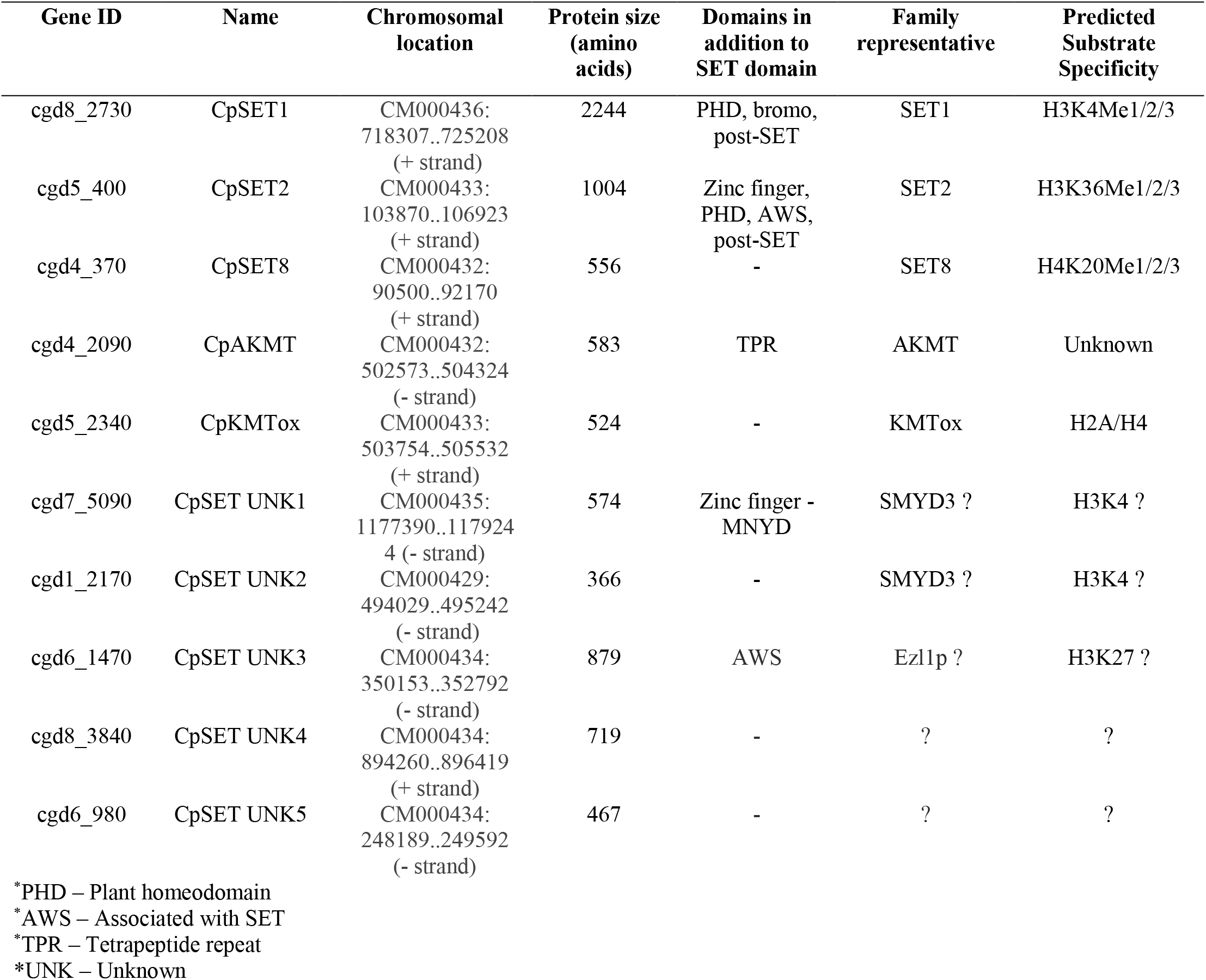
List of putative KMTs identified in the *C. parvum* genome.

The corresponding genes are distributed on 6 chromosomes and the size of the full-length proteins range from 467 to 2244 amino acids. *In silico* analysis also showed the presence of CpSET proteins orthologues in all *Cryptosporidium* species including *C. muris, C. andersoni, C. hominis, C. meleagridis* and *C. ubiquitum*. Among the known methylated lysines, only H3K79 is known to be methylated by DOT1, a non-SET domain histone methyltransferase in mammalians. Unlike SET domain proteins, DOT1 domain containing proteins were observed only in *C. muris* and *C. andersoni*. Analysis of domain organisation of so-called CpKMTs identified additional domains, including the bromodomain, PHD (plant homeodomain) zinc finger domain, associated with SET (AWS) domain, tetratricopeptide repeat (TPR) domain and MYND (<u>My</u>eloid translocation protein 8, <u>N</u>ervy, and <u>D</u>EAF-1). In contrast, JmJC-domain present in lysine demethylases (KDMs) was not identified in *Cryptosporidium* genomes.

The SET domain proteins are usually classified into seven families including SUV39, SET1, SET2, EZ, RIZ, SMYD and SUV4-20 as well as a few orphan members SET7/9 and SET8 (also called as Pr-SET7). Sequences of CpSET domains were aligned with the representatives of these families. Sequence analysis of SET domain of KMTs revealed the presence of four signature motifs namely motif I (GxG), motif II (YxG), motif III (RFINHxCxPN) and motif IV (ELxFDY). As a result, 8 out of 10 predicted CpSETs exhibited at different levels of similarity the three catalytically essential motifs GxG, RFINHxCxPN and ELxFDY of the SET domain (Figure 1) which was not the case for the two remaining CpSETs (cgd6_3840 and cgd6_980), which were therefore excluded from our analysis. The motif II was found to be very well conserved in cgd8_2730, cgd5_400 and cgd4_370 compared to other CpSETs. The sequence between motif II and motif III was observed to be the most variable region of the SET domain. The canonical preSET domain was not reported preceding any of the CpSETs according to InterPro database. However, cysteine (Cys) rich cluster was observed flanking the N-terminal extremity of cgd5_400 and cgd6_1470 SET domains. This region has been identified as AWS domain by InterPro database. Regarding the C-terminal flanking region of the CpSET domain, it is composed of the post-SET domain which contains the CXCX2-4C motif, well conserved in cgd8_2730, cgd5_400, cgd1_2170 and cgd6_1470. This region has also been identified as post-domain in InterPro database. All the CpSETs exhibiting the post-SET domain also conserved the Cys residue in the motif III. Indeed, the Cys residues from post-SET domain and motif III together form a channel to accommodate target lysine side chain. However, cgd7_5090 showed sequence variation in the post-SET motif (CXCX_2_C) similar to SMYD (SET-and MYND-domain containing) and Suv4-20 KMT families. With a distinct residues conserved in motif III, cgd5_2340 is recognized to be part of separate family of KMT called as KMTox. Finally, cgd4_2090 was identified to conserve the post-SET cysteine cluster (CXCX_2_CX_11_CX_2_C) which has been previously described to be a feature of <u>A</u>pical <u>l</u>ysine <u>m</u>ethyl<u>t</u>ransferases (AKMT), a cluster of KMTs only including apicomplexan homologues (Figure 1).

**Figure 1.**
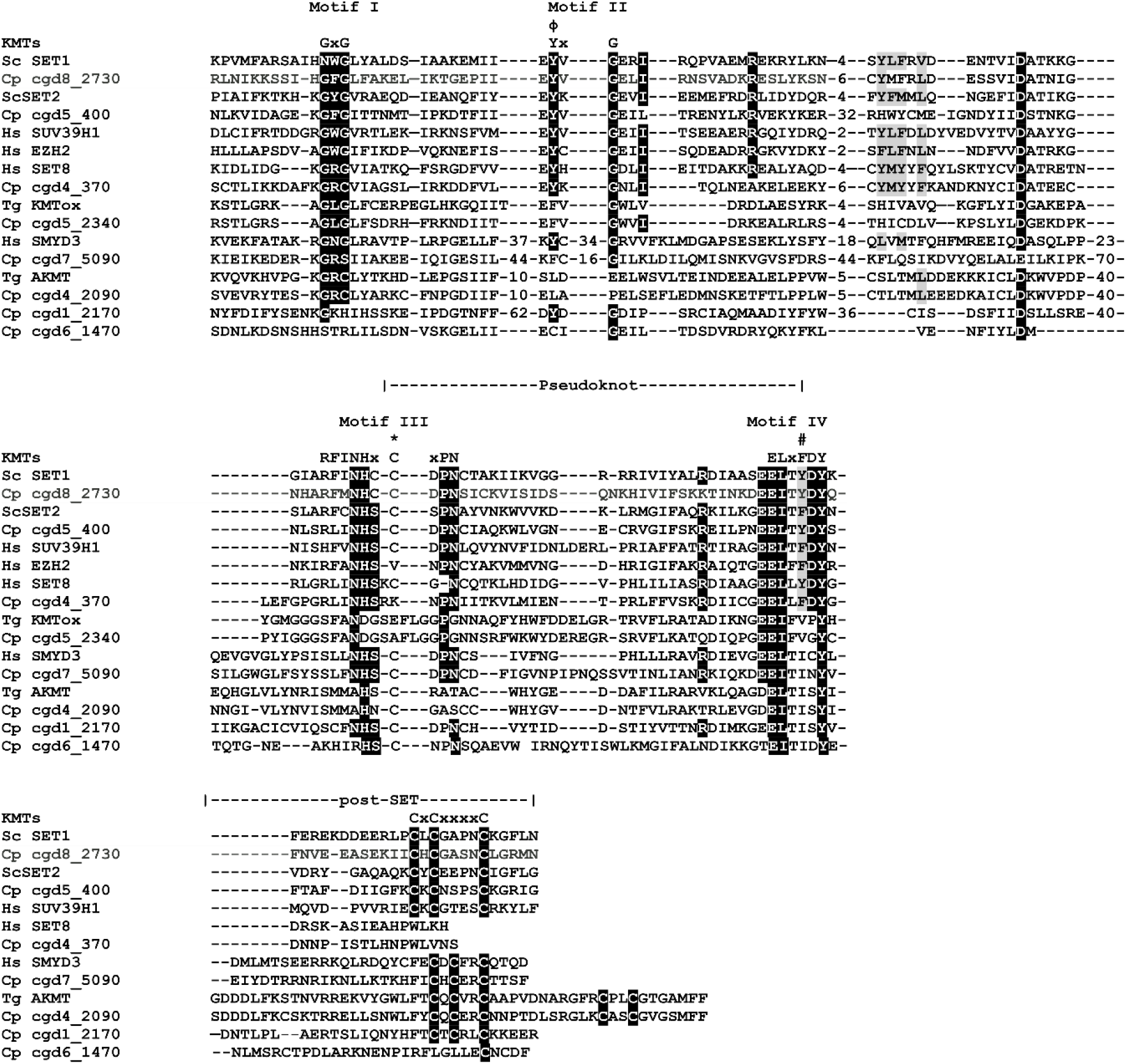
Alignment of SET and post-SET domain sequences of putative CpKMTs with representatives of KMT families. The SET domain is divided into four motifs (I-IV) and their consensus sequences are indicated above the alignment. Motif III and IV are involved in formation of pseudoknot structure to form the active site. The white text on black background indicates identical residues; black text on gray background indicates conserved residues. The residues representing catalytic site are indicated with phi (ϕ). The residues representing F/Y switch are indicated with hash symbol (#). The Cys residue from Motif III that is involved in Zn cluster formation with the post-SET domain are indicated with asterisks (*).

### 2.2. Phylogenetic analysis of putative CpSETs

The 8 *C. parvum* amino acid sequences identified as putative CpSETs were subjected to phylogenetic analysis along with representatives of different substrate specific SET domains (Figure 2). cgd8_2730 clustered with representatives of the SET1 family of HKMTs such as *S. cerevisiae* SET1 and *H. sapiens* SET1 and was thus identified as CpSET1. This distribution was strongly supported by bootstrap resampling in NJ (86%) and ML (97%) methods. The clustering of CpSET1 together with previously identified apicomplexan homologues from *T. gondii, P. falciparum* and *T. annulata* was also moderately supported by bootstrap values (51% and 65% of the replicates under NJ and ML, respectively). The substrate specificity of this family is at the 4^th^ lysine residue of the histone 3 protein (H3K4). Thus, we predicted H3K4 to be the substrate of CpSET1 (Figure 2) (Table 1).

**Figure 2.**
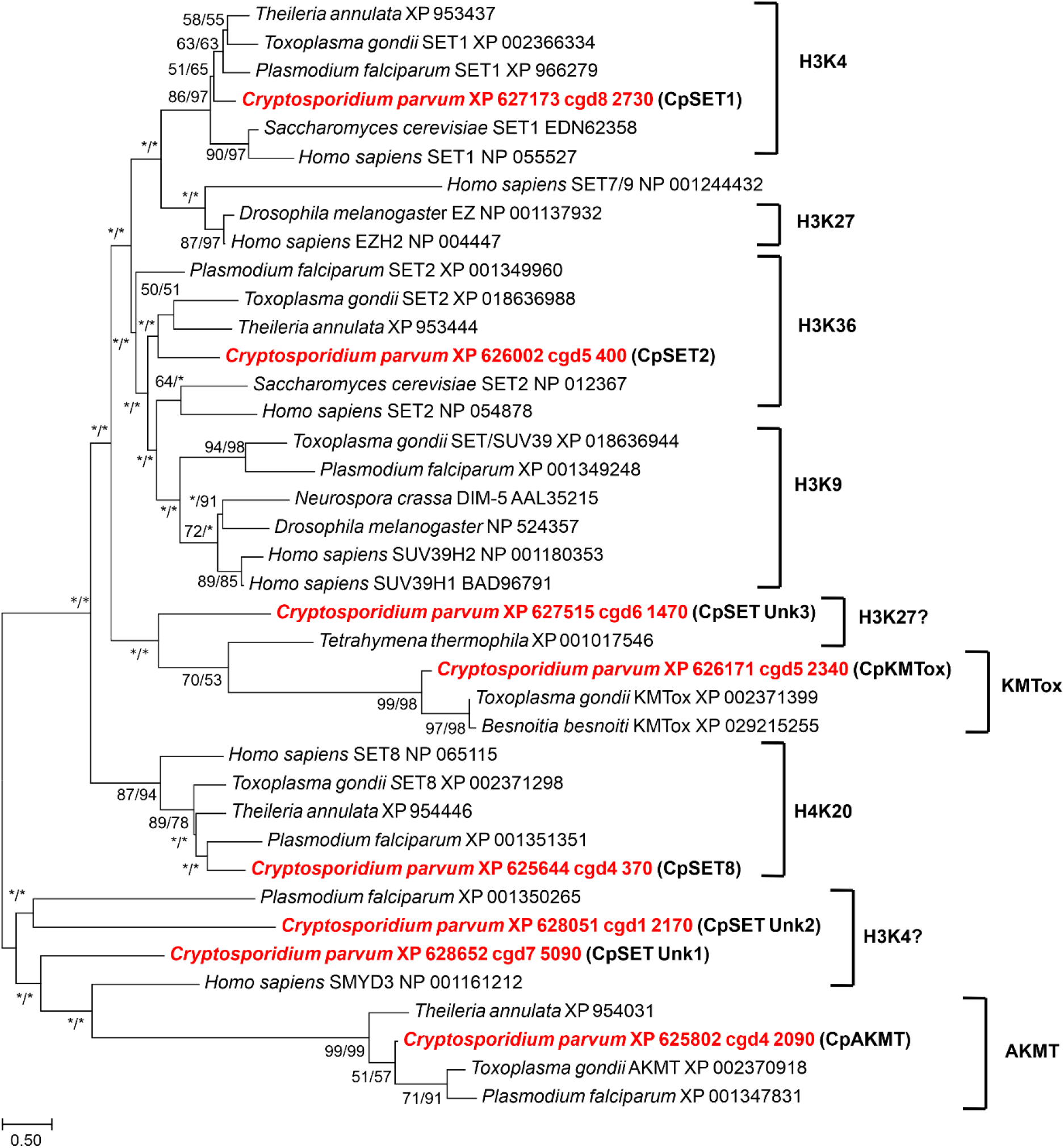
Phylogenetic analysis of SET domain proteins of *C. parvum*. All the putative *Cryptosporidium* sequences are highlighted in red. Numbers near the individual nodes indicate bootstrap values given by NJ (left of the slash) and Maximum Likelihood (right of the slash). Asterisks indicate nodes with bootstrap values below 50%. Branch lengths are proportional to sequence divergence and can be measured relative to the scale bar. The scale bar indicates the branch length corresponding to 0.50 substitutions per site. The predicted substrate specificities of KMTs are also indicated on the right of the figure. The substrate specificities of characterized KMTs clustering with the Cp SET domain proteins are also indicated on the right of the figure. Except KMTox and AKMT are not “substrate specificities”

Regarding cgd5_400, this sequence fell within the paraphyletic SET2 family of KMTs including *Saccharomyces cerevisiae* and *Homo sapiens* as well as apicomplexan enzymes from *T. gondii, T. annulata* and *P. falciparum*. Our phylogenetic tree suggested that cgd5_400 belongs to SET2 family of HKMTs which have been reported to methylate H3K36 and is so-called CpSET2 (Figure 2) (Table 1).

cgd4_370 was observed to be highly homologous to human SET8 and therefore named CpSET8. This homology was strongly supported by bootstrap values of 87% and 94% according to NJ and ML methods, respectively, in our phylogenetic analysis, proposing that CpSET8 methylate H4K20 as described for human SET8. Another putative KMT of *C. parvum* cgd5_2340 grouped together with KMTox from the Apicomplexa *T. gondii* and *Besnoitia besnoiti* with strong bootstrap values of 99% (NJ method) and 98% (ML method) and was thus reported as CpKMTox. KMTox has been defined as a new family of nuclear KMTs specifically found in Apicomplexa and thus including *Cryptosporidium*. Similar to KMTox, AKMTs also form a distinct clade from other known KMTs only found in apicomplexan species. cgd4_2090 or CpAKMT represents *Cryptosporidium* AKMT as it clustered with high bootstrap support (bootstrap values of 99% and 99% according to NJ and ML methods, respectively) with other apicomplexan homologues.

The phylogenetic emergence of the three remaining CpSETs so-called CpSET Unk1 (cgd7_5090), CpSET Unk2 (cgd1_2170) and CpSET Unk3 (cgd6_1470) remained uncertain according to our present study (Table 1). Indeed, CpSET Unk3 branched with unsupported bootstrap support at the base of a large group including KMT of the protozoan ciliate *Tetrahymena* and the KMTox of Apicomplexa. Since the KMT of *Tetrahymena thermophila* is known to methylate H3K27, CpSET Unk3 could be involved in the same methylation event even if no sequence has been yet reported to be associated with H3K27 substrate specificity in Apicomplexa (Figure 2).

The two others CpSET Unk1 and Unk2 are representative of a paraphyletic group that includes in particular the human KMT called SMYD3 known to methylate H3K4. Consequently, the weak grouping of CpSET Unk1 and CpSET Unk2 with SMYD3 speculates a possible methylation of H3K4 by these two KMTs of *Cryptosporidium* (Figure 2).

To complete this overview, unlike other apicomplexan parasites, no *C. parvum* sequence was found to be associated with the SUV39 and EZ families mediating H3K9 and H3K27 methylation, respectively (Figure 2).

### 2.3 Structural characteristics of CpSETs

Homology modelling of the structures of CpSET1, CpSET2 and CpSET8 using publicly available X-ray crystal structures of homologous enzymes showed that the overall architecture of the SET domains belonging to different subfamilies of KMTs were nearly identical. The MolProbity scores evaluating the quality of the models were good with values of 1.62 (92^nd^ percentile), 1.73 (88^th^ percentile) and 1.57 (93^rd^ percentile) for CpSET1, CpSET2 and CpSET8, respectively (Table 2).

**Table 2.** MolProbity statistics for 3D models of CpSETs

| 3D model | Clashscore<sup>a</sup> |  | Ramchandran favoured | MolProbity Score<sup>b</sup> |  |
| --- | --- | --- | --- | --- | --- |
| CpSET1 | 2.24 | 99<sup>th</sup> percentile | 95.04% | 1.62 | 92<sup>nd</sup> percentile |
| CpSET2 | 2.91 | 98<sup>th</sup> percentile | 89.22% | 1.73 | 88<sup>th</sup> percentile |
| CpSET8 | 4.61 | 96<sup>th</sup> percentile | 94.52% | 1.57 | 93<sup>rd</sup> percentile |

<sup>a</sup> percentile score established with N =1784, considering all resolutions
<sup>b</sup> Percentile score established with N=27675, considering structures at all resolution ranges

Pairwise structure comparisons with DALI program was used to check whether conserved residues line up between CpSETs and the templates. As described in Table 3, CpSET1 shares highest structural identity with the templates (52%) compared to CpSET2 (43%) and CpSET8 (44%). The visualization of these superimposed structures was performed using ChimeraX software (Figure 3).

**Table 3.** Pairwise structure comparison between generated models and templates.

| **Structure**<br>**Parameters** | **CpSET1 onto 5F6L** | **CpSET2 onto 6J9J** | **CpSET8 onto 5TEG** |
| --- | --- | --- | --- |
| RMSD (Å)<sup>a</sup> | 0.3 | 0.6 | 0.4 |
| Number of<br>superimposed residues | 136 | 139 | 144 |
| Total number of<br>residues | 143 | 169 | 148 |
| Dali z-score<sup>b</sup> | 24.4 | 22.2 | 25.8 |
| Percentage identity | 52 % | 43 % | 44 % |

<sup>a</sup> RMSD is the root-mean-square deviation computed over the Cα atoms of superimposed residues.
<sup>b</sup> A DALI z-score greater than 3 is considered as reflecting similar structures. Higher z-scores indicate more similar structures.
<sup>c</sup> the percentage identity is calculated between structurally superimposed residues.

**Figure 3.**
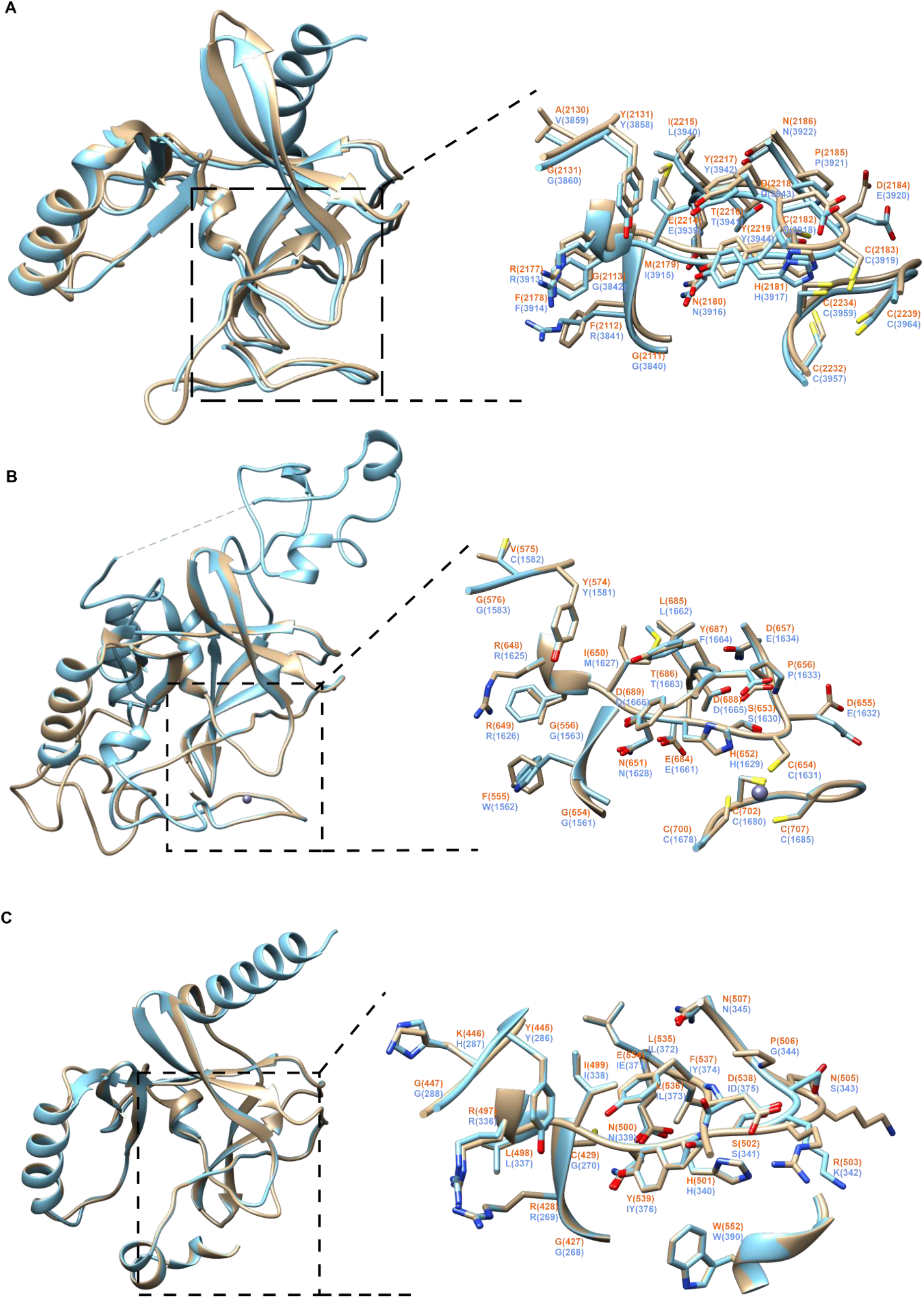
Structural modelling of SET domain regions of *C. parvum* KMTs. Superimposition of 3D homology models of SET domains of [sandy brown] CpSET1 (A), CpSET2 (B) and CpSET8 (C), coloured in sandy brown, with, in cyan, those of resolved crystal structures of SET domains of *Homo sapiens*, HsSET1, HsSET2 and HsSET8, respectively. The superimposition of the catalytic site has been highlighted in the black dotted box and depicted in an enlarged view with side chains showed in stick representation (nitrogen and oxygen atoms in blue and red, respectively) and labelled in orange and blue for *C. parvum* and *H. sapiens* structures, respectively.

Briefly, all the structures contained the specific β fold identified only in KMTs, not in any other previously characterized AdoMet-dependent methyltransferases. The fold has several series of curved β strands forming several small sheets that define the core of the SET domain. This β fold is followed by knot-like structure which is also observed in all the 3D models of CpSETs. The knot involves a C-terminal β strand threading through a hoop consisting of two β strands and their connecting region. This represents an archetypal feature of SET domain which consists of residues from the motif II, motif III and post-SET region that enclose the lysine residue and holds it in the appropriate chemical environment and position for methyl transfer by motif I (Figure 3, boxed in black dotted line). The essential residues have similar arrangement in all the SET domains (Figure 3, enlarged images of active site).

CpSET1 (Tyr 2129 and Tyr 2217) and CpSET2 (Tyr 574 and Tyr 659) conserved the key tyrosine residues (Figure 3A; 3B). These Tyr residue are expected to form an intricate network of hydrogen bonds which would place the methyl group in direct line with the N_ε_ of the lysine residue of the histone tail. Structural alignment revealed that in CpSET8 one of these tyrosine residues is replaced by phenylalanine residue (Tyr 445 and Phe 537) (Figure 3C). This represents the F/Y switch which determines the KMT can mono-, di- or tri-methylate the histone.

The C-terminal flanking region of different SET domain families is often divergent but CpSET1 and CpSET2 exhibited a classical post-SET domain. The prominent feature of this domain is a zinc-binding cage formed by three Cys residues from the C-terminal region whereas the fourth tetrahedral Cys ligand (CpSET1 Cys 2183; CpSET2 Cys 654) is provided by the loop linking motif II and motif III of SET domain. The narrow channel formed as a result of Cys interactions accommodates the target lysine and brings the Nε in close proximity of the donor at the opposite end of the channel (Figure 3A, B). In CpSET8, the C-flanking domain consists of a helix and the presence of the Trp 552 residue is responsible for interactions with the cofactor (Figure 3C).

### 2.4 Enzymatic activity of CpSET8

To check whether CpSETs represent active KMTs, we expressed and produced SET domain regions of CpSET1 and CpSET8 in a bacterial system. Western blots analysis showed detectable amounts of CpSET8 in the soluble fraction after induction and lysis of bacteria compared to CpSET1 (Figure 4A). Therefore, KMT activity assay was performed on the induced soluble protein fraction of bacterial lysate containing SET8. As a result, CpSET8 showed HKMT activity against H4 which evidenced by a relatively strong band seen below 15kDa (Figure 4B, Lane 2) compared to the controls which did not contain either SAM (Figure 4B, Lane 1) or H4 (Figure 4B, Lane 3). The specificity of H4K20 methylation band was also confirmed by the detection of H4 at the same molecular weight. Moreover, since anti-his tag antibody detects the 6X histidine tag present in the recombinant CpSET8 and, equal intensity of anti-his tag bands corresponds to equal amount of protein loaded in each well, this indicated that the high intensity band detected (red arrow) was the result of enzymatic activity of CpSET8.

**Figure 4.**
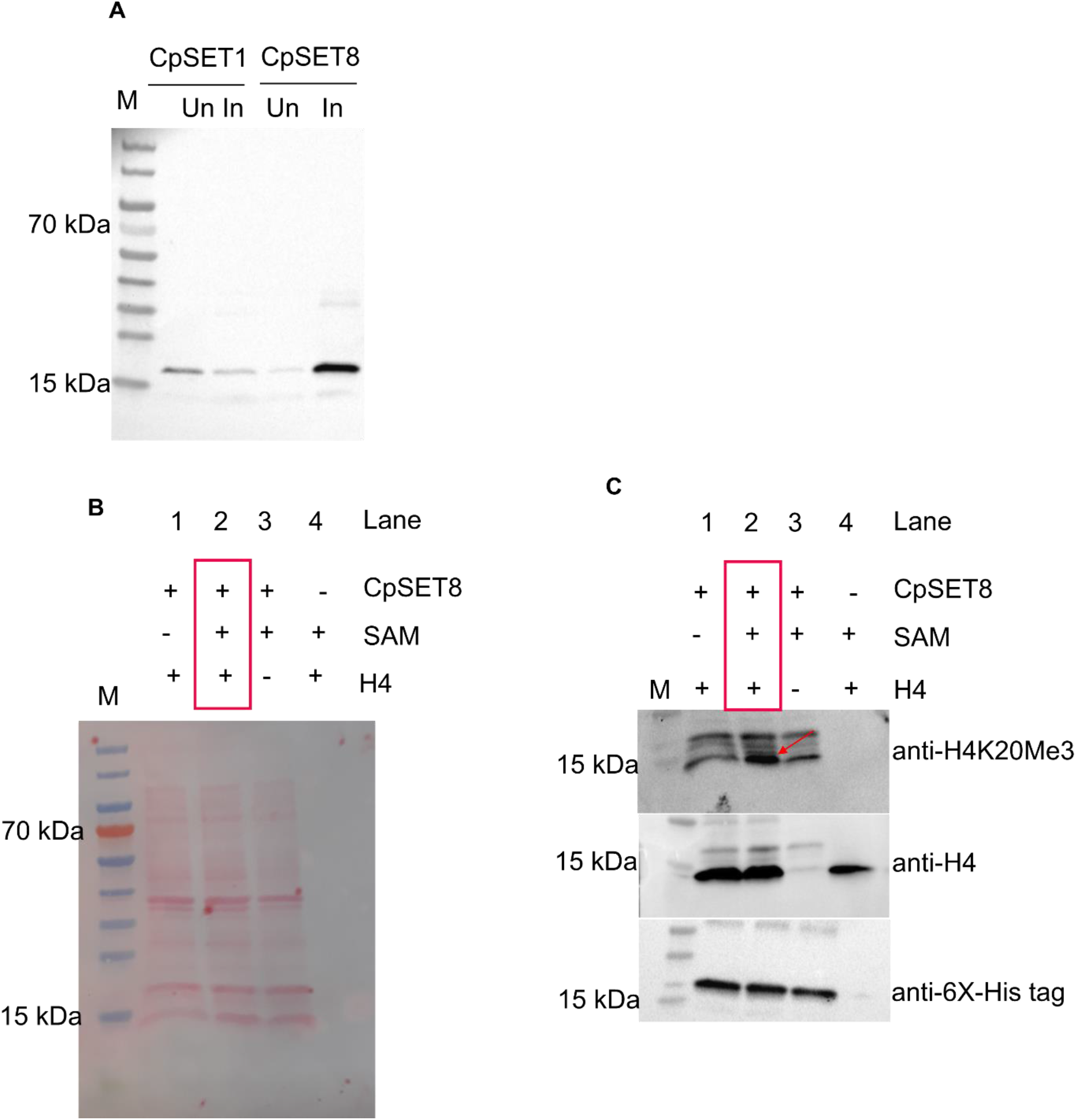
Expression of putative CpSETs and enzymatic activity of CpSET8. A. Western blot analysis of 6x-histidine tagged CpSET domains using bacterial expression system. Anti-6x-His tag antibody was used to detect the CpSETs. M – Molecular marker, Un – Uninduced soluble protein fraction, In – Induced soluble fraction. B. Ponceau S staining of the western blot subjected to chemiluminescent detection of H4 methylation by recombinant 6x-histidine tagged CpSET8. C. Chemiluminescent signals showing CpSET8 to tri-methylate H4 using anti-H4K20Me3 antibody (red arrow). Recombinant human H4 was detected using anti-H4 antibody. CpSET8 was detected using anti-6x-His tag antibody. H4 – Histone 4. SAM – S-adenosyl methionine.

### 2.5 Functional characterization of identified KMTs during *C. parvum* life cycle

To determine the expression pattern of KMTs during the *C. parvum* biological cycle *in vitro*, RT-qPCR analysis of the eight *CpSET* genes was performed. The expression of all *CpSET* genes was determined at 3 time points i.e. 6 h, 24 h and 55 h PI, relative to their expression at 2 h PI. *CpSET1* showed 14-fold increase at 6 h PI, corresponding to the time at which trophozoite development is predominant. Thus, *CpSET1* is the highest expressed KMT gene, particularly during initial stages of the infection. *CpAKMT* expression was 10-fold increased during meront development at 24 h PI. *CpSET2*, *CpSET8*, and *CpKMTox* genes also showed high expression during trophozoite stage (6 h PI) followed by asexual (24 h PI) and sexual stages (55 h PI). At 55 h PI, wherein the sexual stages were predominant, all the identified putative *CpKMTs* were constitutively expressed. The gene expression of putative unknown *CpKMTs* such as *CpSET Unk1*, *CpSET Unk2* and *CpSET Unk3* during intracellular development was observed to be lower compared to that observed during the extracellular sporozoite stage at 2 h PI (Figure 5A). Out of the five reported putative CpKMTs, CpSET1, CpSET2 and CpSET8 have been identified as histone lysine methyltransferases (HKMTs). In order to functionally characterize these HKMTs, the histone lysine methylation events associated with their activity during the life cycle of *C. parvum* were investigated *in vitro* in relation with the different parasite developmental stages. As previously described, CpSET1, CpSET2 and CpSET8 are associated with H3K4, H3K36 and H4K20 methylation. Immunofluorescence analysis revealed that H3K4, H3K36 and H4K20 methylations occur during all the intracellular stages of the parasite. It can be observed that anti-tri-methyl H3K4 (H3K4Me3) antibody recognized the *C. parvum* chromatin with a broader nuclear distribution through all the developmental stages. Regarding anti-tri-methyl H3K36 (H3K36Me3) and anti-tri-methyl H4K20 (H4K20Me3) antibodies staining, it showed a punctate pattern spread throughout the nucleus, most likely the pericentric heterochromatin during the intracellular stages compared to extracellular sporozoite stage (Figure 5B). By quantifying the labelling of these histone lysine methylations by western blotting, we were able to confirm that H3K4 methylation levels remain consistent during meronts development at 24 h PI and microgamonts and macrogamonts development at 55 h PI. On the other hand, H3K36 and H4K20 methylations levels have been shown to fluctuate from asexual (24 h PI) to sexual stage 55 h PI (Supplementary Figure 2A). H3K36 and H4K20 methylations increase by 2 and 3 fold respectively, when there is predominance of sexual stages of the parasite in the culture (Supplementary Figure 2B).

**Figure 5.**
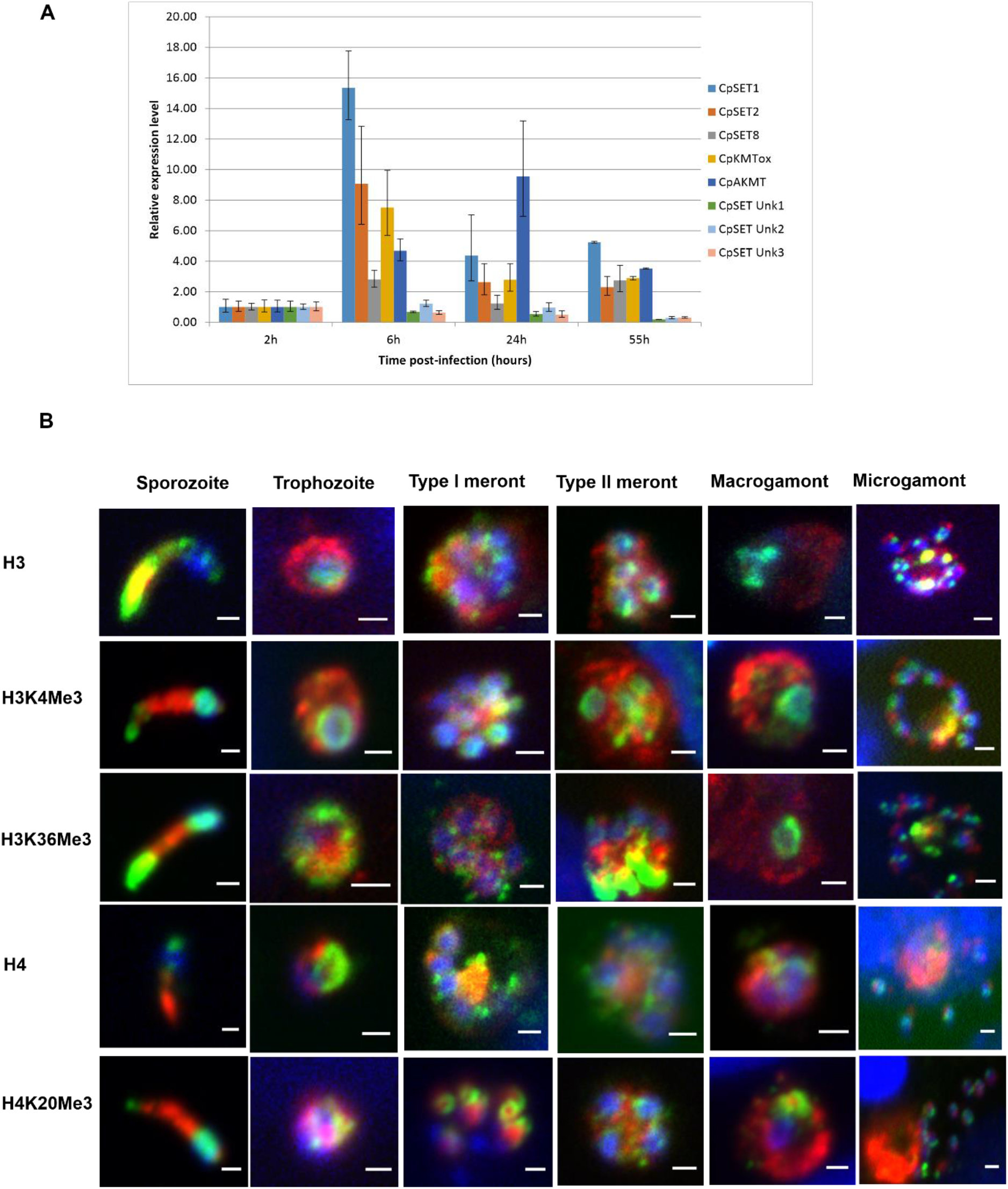
Characterization of KMTs during *C. parvum* infection in *in vitro* culture. A. RT qPCR analysis illustrating expression of *CpSET* genes during *C. parvum* development *in vitro*. The expression levels were analysed in triplicates and normalized with 18S rRNA gene as internal control. The ΔC_t_ values at the sporozoite stage were used as the calibrator B. Immunofluorescence analysis of histone lysine modifications in different stages of *C. parvum*. Co-staining with anti-histone antibodies such as H3 and H4 and anti-histone methylation antibodies such as H3K4Me3, H3K36Me3 and H4K20Me3 (green), anti-*Cryptosporidium* antibody (red) and DAPI (blue).

### 2.6 Effect of *C. parvum* infection on methylation of the host histone lysine proteins

The well-documented model of *C. parvum*-induced colon cancer in SCID mice treated with dexamethasone was used. The detection of *C. parvum* infection in these animals was confirmed by quantification of the oocyst shedding for the entire duration of the experiment. Upon histological examination of the ileo-caecal region of infected animals, the presence of well-differentiated adenocarcinomas invading the submucosae through the *muscularis mucosae* was confirmed after 60 days PI. The availability of antibodies to detect histone lysine methylations in the parasite allowed us to decipher the effect of *C. parvum* infection on the host epigenome. Immunofluorescence analysis of H3K9 revealed a strong immunofluorescence signal in the epithelium of the ileo-caecal region of *C. parvum* infected animals at 60 days PI, while a significant decrease in the H3K4, H3K36, H3K27 and H4K20 methylations was observed. H3K18 methylation levels were shown to be unaffected (Figure 6A). The endogenous levels of H3 protein were used as control. Immunofluorescence staining revealed strong signals in parasite nuclei with all antibodies except H3K18. Quantification of fluorescence intensity signals is available in supplementary figure 3. Further, western blotting analysis was performed to determine which stages of the parasite could affect the host methylation events during *C. parvum* infection *in vitro*. As a result, we observed downregulation of all the methylation marks except for H3K18 to be downregulated by 0.5 fold in infected HCT-8 cells relative to non-infected HCT-8 cells at 55h PI (Figure 6B; supplementary Figure 4C). H3K36 and H4K20 methylations were also relatively downregulated by 0.5 fold in infected host cells during the peak of meront stage development of the parasite at 24h PI (Figure 6B, supplementary figure 4C). H3K4, H3K36, H3K27 and H3K9 methylations were relatively upregulated in the host during the trophozoite stage development of the parasite at 6h PI (Figure 6B, supplementary figure 4C).

**Figure 6.**
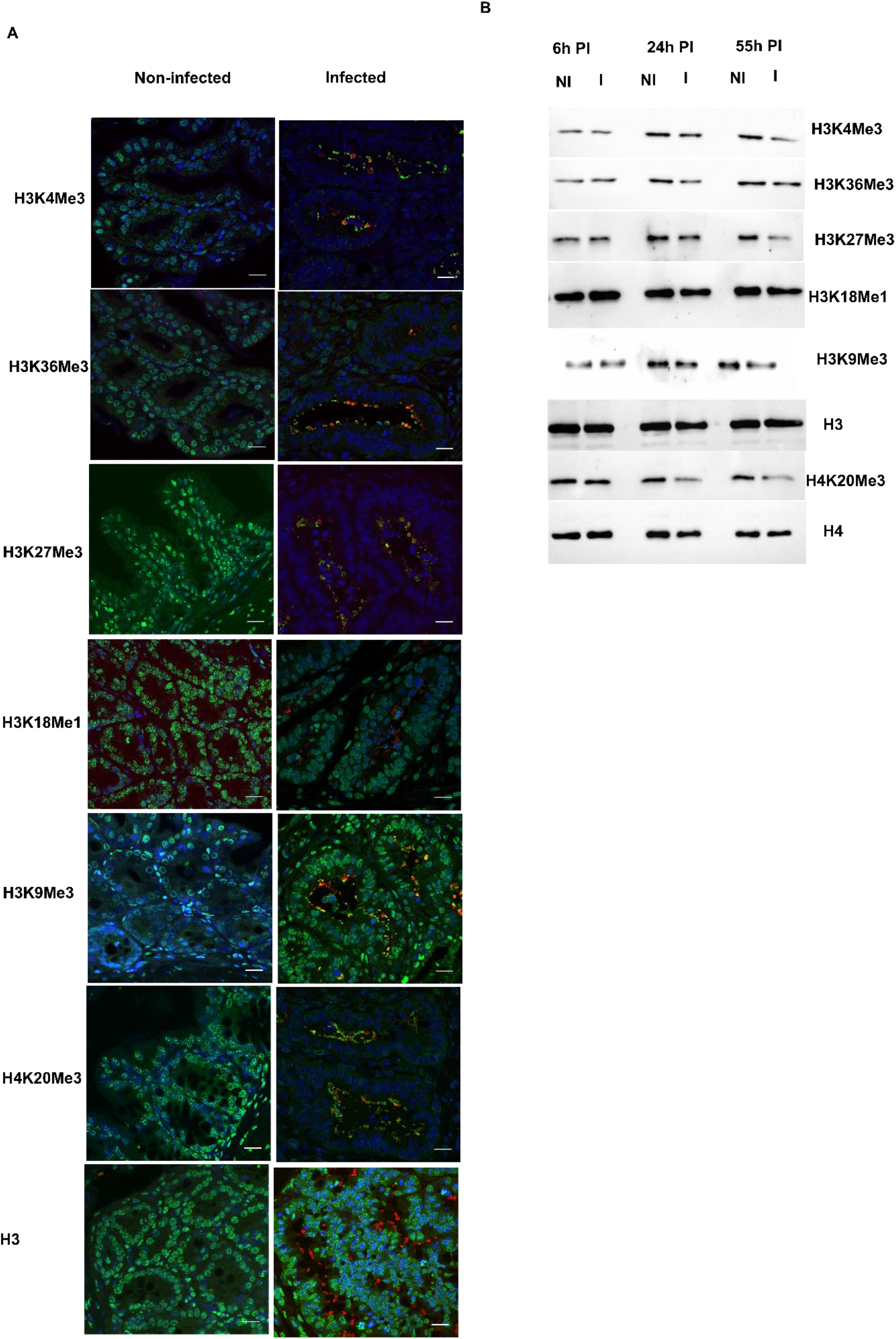
Effect of *C. parvum* infection on the host methylation epigenome. A) Immunofluorescence analysis of histone methylation events during *C. parvum* infection *in vivo* at day 60 PI. Co-staining with anti-histone methylation antibodies such as H3K4Me3, H3K36Me3, H3K27Me3, H3K18Me1 and H4K20Me3 (green), *anti-Cryptosporidium* antibody (red) and DAPI (blue) of the ileo-caecal region of *C. parvum* infected SCID mice. The difference in the fluorescence intensity was observed to be significant (p<0.05). B) Western blotting analysis of histone methylation events during *C. parvum* infection *in vitro* at 6 h PI (trophozoite stage), 24 h PI (asexual stage), 55 h PI (sexual stages) after purification of histones from host cell. NI – Non-infected HCT8 cells. I – Infected HCT8 cells. Scale bar – 20 μm. Results are representative of three independent experiments.

## 4. Discussion

In the current study we investigated the potential significance of an epigenetic modification known as histone lysine methylation during *C. parvum* infection. Identification of structurally active KMTs followed by their functional characterization in the form of lysine methylation events makes this study the first one to recognize the importance of epigenetics in the development and potential pathogenicity of *Cryptosporidium*.

Several proteins including KMTs which are responsible for the methylation of specific residues have been characterized in higher eukaryotes, and all but one of these enzymes possess a SET domain. A first screening using InterPro database identified 10 SET-domain containing proteins in *C. parvum* so-called CpSET proteins. A further analysis emphasized their conservation in all other sequenced *Cryptosporidium* genomes suggesting the parasite ancestor had acquired these genes before speciation and divergence within this genus. However, KMTs belonging to the DOT1 family were detected in *C. muris* and *C. andersoni* but not in *C. parvum*. The absence of DOT1 domain containing lysine methyltransferases in some *Cryptosporidium* species as in related genera including *Toxoplasma* (32) and *Plasmodium* (27) suggests a secondary loss of the corresponding genes during the evolution of the phylum Apicomplexa. Along with the SET domain, the presence of PHD zinc finger domain (33) and bromodomain (34) regards the CpSETs proteins as capable of forming protein complexes and interact with the chromatin. Therefore, it can be suspected that PHD zinc finger domain and bromodomain facilitates binding of the CpSET proteins chromatin and assist in lysine methylation of histones which is known to be catalyzed by SET domain.

The histone modifications catalyzed by KMTs possess narrow substrate specificities, often targeting a single lysine within the respective substrates. These KMTs also differ in their preference for different methylation states (mono-, di-, or tri-methylation) of lysine residues. In spite of the conserved overall structural plasticity, the variations at the active sites have been identified to contribute to their varying substrate specificities (35). Hence, we decided to align the primary sequence of the SET domain of identified CpSET proteins with different families of known KMTs. The SET domain consists of four signature motifs; motif I to IV belong to the pre-SET region of the SET domain, out of which the motifs III and IV are highly conserved and form a pseudoknot structure. The structural significance of this knot is to bring two conserved motifs together in order to form an active site immediately close to the motif I which is a binding site for the methyl donor S-adenosylmethionine (AdoMet). Thus, identification of these catalytically essential motifs involved in transfer of methyl group in 8 out of 10 putative CpSETs allows speculating that *C. parvum* KMTs are functionally active.

Out of all the putative CpKMTs, CpSET1, CpSET2 and CpSET8 were identified as potential HKMTs. Especially, all the residues of the signature motifs of SET and post-SET domains were conserved in the CpSET1 primary amino acid sequence. Moreover, CpSET1 clustered together with other SET1 family homologues with high bootstrap support in the present phylogenetic analysis. ScSET1, one of the homologues grouping together with apicomplexan enzymes has been associated with tri-methylation of H3K4 in the regions of genes which are transcribed early, thus, making them a mark of transcription activation (36). Moreover, domain organization of CpSET1 consists of bromodomain and PHD domain which have been reported to interact with trimethylated H3K4 (37), suggesting that CpSET1 employs these interacting domains to target euchromatin and methylate H3K4. In addition, superimposition of CpSET1 3-D model with the available crystal structure of human MLL1, one of the members of the SET1 family, strongly suggest that CpSET1 can be considered as a structurally active HKMT. MLL1 has been shown to catalyze multiple methylations. The superimposition study has shown that CpSET1 conserves all the active site residues (Phe 2159, Tyr 2217, Tyr 2219, Phe 2221, Cys 2156 to Phe 2158) found in SET domain of MLL1. Moreover, MLL1 SET domain has similar arrangement to other SET domains (38). CpSET1 also conserved key tyrosine residues (Tyr 2129 Tyr 2217) required for the transfer of methyl group. Even though, sequence analysis indicates that these tyrosine residues constrain HKMTs such as SET8 to be mono-methylases. It can be speculated that certain modulations in the configuration of the active site of CpSET1 could allow multiple methylations which is the case for MLL1. Superposed structure of MLL1 SET domain with other HKMTs (SET7/9, SET8 and Dim5) revealed that MLL1 has a more spacious active site (38). This spacious active site is attributed to shift in the orientation of SET-I region and C-terminal flanking region in MLL1. This feature is also evident in CpSET1 after superimposition with MLL1. The CpSET1 residues (Cys 2156 to Phe 2158) are conserved which are responsible for this change in orientation and allowing free movement of the lysine side chain. Thus, CpSET1 can be predicted to mono, di or tri-methylate H3K4. Interestingly, MLL family members (MLL1-4, SET1A and SET1B) are known to methylate H3K4 and have pivotal roles in the regulation of the transcription of genes involved in development and hematopoiesis (38). In particular, cell cycle progression (39). In this context, the immunofluorescence labeling observed during the development of the parasite using anti-H3K4Me3 antibody could be attributed to the existence of a functional CpSET1 representing a MLL1 member of the HKMT family *Cryptosporidium*. Moreover, the consistent high expression of *CpSET1* gene during the parasite development suggests that H3K4 methylation is not a dynamic but a rather stable post-translational modification necessary during all the stages of the parasite life cycle. In addition, H3K4Me3 has been shown to mark the promoter of actively transcribed genes in different apicomplexan parasites such as *T. gondi* (40), *T. annulata* (41) and *P. falciparum* (25). Hence, the stable nature of this mark can be attributed to the fact that this modification is involved in expression of actively transcribing genes during each stage of the parasite. However, the open conformation of the SET domain of MLL1 has been predicted to be not efficient to transfer the methyl group from the cofactor S-adenosyl-L-methionine (AdoMet) to the target lysine (38). Recently, other components such as RBBP5-ASH2L have been identified to bind and activate MLL family methyltransferases through a conserved mechanism (42). As the SET domain of CpSET1 is structurally identical to MLL1 SET domain, the involvement of other components needed by CpSET1 to maintain its HKMT activity has to be investigated in further studies. Even though, we made preliminary attempts to clone and purify the CpSET1 domain in a bacterial system, western blots analysis did not show detectable amounts of CpSET1 in the soluble fraction after induction and lysis of bacteria. Further studies should work on standardizing the solubility protocols for the concerned domain.

CpSET2 and CpSET8 were the other two HKMTs of *Cryptosporidium* identified to be structurally active. Even though, the phylogenetic tree reconstructed in the present study do not strongly support the grouping of CpSET2 with homologous enzymes from Apicomplexa and others eukaryotes, analysis of its primary sequence revealed the presence of a typical post-SET domain and a cysteine rich N-terminal region preceding the SET domain which both represent peculiarities of proteins belonging to the SET2 family of HKMTs (43). Moreover, the superimposition of 3-D homology model of SET domain of CpSET2 with a template SET domain of SETD2 (protein data bank: 6J9J.A) revealed 43 % identity between the two structures. Interestingly, SETD2, is the only HKMT known to mediate tri-methylation of H3K36, whereas other H3K36 methyltransferases can only mono or di-methylate H3.

Strikingly, H3K36me3 is known to be mediated by a single HKMT such as SETD2, whereas other H3K36 methyltransferases can only mono- and di-methylate H3K36 (44). Intriguingly, mutating arginine residue (Arg1625Cys) present in the SET domain results in an enzymatically inactive SETD2 failing to tri-methylate H3K36 (45). This arginine residue (Arg 648) is conserved in CpSET2. Thus, based on the structurally conserved residue it can be predicted to tri-methylate H3K36 in *Cryptosporidium*. In addition, the use of anti-H3K36Me3 antibody, detected this methylation mark to be distributed throughout the life cycle stages of the parasite. Thus, functionally validating the speculation of active CpSET2 to be present in *Cryptosporidium*. However, H3K36Me3 has been reported to be a mark of gene activation in higher eukaryotes (46). But methylation of H3K36 has been linked to repression of *var* genes in *Plasmodium* parasites (Jiang L et al. 2013). In contrast, *Theileria* genome did not show any regions of strikingly high H3K36Me3 enrichment (41). Thus, the significance of this methylation mark in *Cryptosporidium* remains to be explored.

SET8 family of HKMTs, have been characterized to mono-methylate H4 in humans (47). The identification and characterization of SET8 related homologs in Apicomplexa such as *Plasmodium* (27) and *Toxoplasma* (48), overruled the thought that this enzyme is restricted to metazoans (35). CpSET8 identified in our study was also predicted to methylate H4K20 according to our phylogenetic analysis. Thus, reinforcing the importance of histone methylations in chromatin structure and function in Apicomplexa. The superimposition of 3-D homology model of SET domain of CpSET8 with a template SET domain of SET8 (protein data bank: 5teg.A) revealed 44 % identity between the two structures. The conserved tyrosine residues (Tyr 245 and Tyr 334) within the active site of human SET8 are responsible for maintaining an intricate network of hydrogen bonds to position the side chain of only mono-methylated lysine residue (47). On the contrary, Dim-5, a HKMT which can tri-methylate its target lysine can accommodate mono-, di- and tri-methylated lysine in its active site. This characteristic of Dim-5 has been attributed to the replacement of one of the tyrosine residues to phenylalanine residue (Tyr 178 and Phe 281) (49). Interestingly, structural alignment revealed that in CpSET8 as well one of these tyrosine residues is replaced by phenylalanine residue (Tyr 445 and Phe 537). Thus, suggesting that CpSET8 is capable of adding multiple methyl group to its target lysine.

Thus, based on the analysis of structural elements, it can be hypothesized that CpSET8 is capable to methylate H4K20Me1, H4K20Me2 and H4K20Me3 like SET8 of *T gondi* (48). This structural analysis was confirmed herein by showcasing HKMT activity of CpSET8 domain for H4K20Me3. Along with the suspected dynamic nature of CpSET8, the methylation events predicted for this HKMT have also been observed to be regulated during parasite differentiation. This phenomenon can be considered as a characteristic of apicomplexan parasites as H4K20Me3 mark in *T. gondii* has been reported to be cell cycle dependent (48). Even then, the possibility of CpSET8 methylating other targets cannot be excluded. However, unlike other HKMTs such as SMYD3 which are known to have multiple targets (50,51). SET8 family of HKMTs have till date only been shown to have a single target that is H4K20 (43). Hence we strongly propose that CpSET8 is capable of methylating H4K20. Indeed, considering these as preliminary results, further studies need to be carried out by mutating the predicted catalytic residues of the whole protein to see if the protein folding has any effect on the function of the SET domain

CpAKMT was another identified KMT which differed in the C terminal region wherein it retains two extra cysteines in addition to the post-SET domain. Phylogenetic analysis revealed that CpAKMT and its homologues from other Apicomplexa clustered together and represented a sister-group of HsSMYD3, as described in previous evolutionary studies including *Plasmodium* (27) and *Toxoplasma* KMTs (52). The lack of MYND zinc finger domain, a fundamental feature of SMYDs allows to group the AKMT homologues as a distinct family of KMTs. Structural comparison between the AKMT of *T. gondii* and SMYD proteins revealed that SMYD1-3 do not seem to possess the necessary features that are compatible for dimer formation whereas AKMT are dimeric enzymes (52). In *T. gondii*, a functional AKMT has been reported to be localized at the apical complex and associated with parasite motility and egress (53). Moreover, lack of a MYND domain in AKMT might correlate with its specific function outside the nucleus (54). Interestingly, immunofluorescence observations detected the labeling of histone lysine methylations at the apical region of *C. parvum* sporozoites and merozoites. In addition, the *CpAKMT* gene had a relatively high expression during the merozoite development and egress in *C. parvum*. In *P. falciparum*, histones released from the parasite exert a disruptive effect on the endothelial barrier function through a charge-related mechanism in order to induce pro-inflammatory responses (55). Along with the extra-nuclear localization of histones, histone modification such as H3K9Me1 in *P. falciparum* has been recognized as a new function linked to the parasitophorous vacuole and the interaction of the parasite with the host (56). Therefore, we hypothesize that CpAKMT is a functional KMT localized at the apical region capable to methylate extra-nuclear histones of *C. parvum*. However, possibility of CpAKMT to methylate non-histone proteins at the apical region to assist in motility cannot be overlooked. Similar to SMYD3 which has been recently characterized as a non-histone methyltransferase considering its cytoplasmic localization (57). Further studies have to be carried out to address this functional aspect.

KMTox was another identified KMT in *C. parvum*. First identified in *T. gondii*, the presence of High Mobility Group (HMG) domain in KMTox allows it to recognize the bent DNA with the SET domain involved in the methylation of histones H4 and H2A *in vitro* (58). Even if the HMG domain was not identified in CpKMTox, the phylogenetic analysis based on SET domain sequences clustered together all apicomplexan KMTox with a high bootstrap value suggesting that CpKMTox could be predicted as a novel histone H4- and H2A-specific methyltransferase. Even though it was previously reported that TgKMTox formed a distinct clade with no obvious homologues in the Apicomplexa lineage (58), our phylogenetic analysis identified other apicomplexan parasites retaining KMTox such as *C. parvum* and *B. besnoiti* that could represent a new clade of KMTs only found in this group of protozoa.

The nine cysteines residues of the pre-SET domain usually found in the SUV39 family of KMTs (59) were not identified in CpSETs. In parallel, none of the CpSETs clustered in our phylogenetic tree with the known homologues belonging to this family as previously reported (24). Thus, it can be speculated that either H3K9 methylation is not a crucial post-translational modification required for the survival of *C. parvum* or that a yet unknown CpSET may carry out this function. The post-SET region of KMTs also consists of conserved cysteine residues (CXCX4C). These three cysteines coordinate a zinc ion tetrahedrally together with cysteine of motif III of SET domain to form a narrow channel to accommodate the target lysine side chain. Consequently, the post-SET region is extremely crucial for the activity of a functional KMT. We observed that in CpSET1, CpSET2, CpSET Unk2 and CpSET Unk3 the cysteines of the post-SET region along with cysteine of motif III were conserved. However, CpSET Unk2 and CpSET Unk3 lacked the presence of the signature of motif I and could not be grouped together with known families of KMTs. On the other hand, CpSET Unk1 presented a variant of post-SET domain which is observed in the SMYD or SUV4-20 families of KMTs. The presence of a MYND zinc finger motif in CpSET Unk1 and its weak association with *H. sapiens* SMYD3 in our phylogenetic tree suggests that this CpSET could belong to the SMYD subfamily, which is composed of H3K4-specific methylase (50). However, due to their high variability in terms of primary sequence, the functions associated with these CpSETs cannot be affirmatively predicted and thus these enzymes were considered as unknowns. Unlike the other characterized KMTs based on phylogenetic and structural analysis which show dynamic expression during the intracellular stages of the parasite, *CpSET Unk1, CpSET Unk 2* and *CpSET Unk 3* expression levels were relatively unaffected. Thus, these putative KMTs are predicted to not play a role in parasite development.

To complete this overview, KDMs containing JmjC-domain were not identified in *Cryptosporidium*, highlighting that mechanism for histone demethylation is not present in this parasite as reported in previous studies (27,60).

Strikingly, new insights into the effect of *C. parvum* infection on the host histone lysine methylation events were provided in the present study. Indeed, the modulation of different lysine methylation marks in *C. parvum* infected mice was described with the upregulation of H3K9me3 and the downregulation of H3K27Me3, H4K20Me3, H3K4Me3 and H3K36Me3. Till date, one single study identified *C. parvum* infection to induce transcriptional suppression of host genes by upregulation of H3K9Me3 methylation (61). In addition, H3K9Me3 and H3K4Me3 are considered to be opposing and mutually exclusive chromatin modifications (62). Accordingly, it can be proposed that during *C. parvum* infection, H3K9Me3 and H3K4Me3 are functionally connected. Interestingly, the neoplasia developed as a result of chronic infection has been speculated to induce massive loss in the methylation levels. The PI3K/AKT signaling pathway known to be activated during *Cryptosporidium* infection (63) can downregulate H3K27 and H3K4 methylations and consequently activate Epithelial Mesenchymal Transition (EMT) in gastric cancers (64) explaining in part the transformation process that takes place in *Cryptosporidium* infected epithelial cells. This result is consistent with a recent microarray study reporting that EMT takes place within the tumor microenvironment induced by *C. parvum* infection in a rodent model (65). Moreover, *in vitro* analysis, revealed that both these methylations decreased over time predominantly during the development of sexual stages. Down regulation of H3K4Me3 mark has been reported on the promoter ofNF-κB-related pro-inflammatory genes in *Leishmania* infected macrophages as a survival strategy of the parasite (66). Unlike *C. parvum, T. gondii* infection induces upregulation of H3K27Me3 mark. This upregulated level of methylation represses pro-inflammatory cytokines genes to allow the persistence of the infection (67). Thus, regulation of histone lysine methylations levels could be one of the immune escape strategies of *C. parvum*. Particularly, during the development of the sexual stage, the parasite can be expected to target the host’s innate immune response genes. In fact, through recent transcriptomics analysis, persistent *C. parvum* infection has been reported to evade host innate immune response to develop tumor microenvironment (65). Further studies have to be conducted to better elucidate this aspect

In conclusion, lysine methylases were successfully characterized in *C. parvum*. Since these enzymes are involved in the development of the parasite life cycle, they may be considered as potential targets to curb the infection opening new avenues for anti-parasite drug discovery. On the other hand, our work contributes to the growing evidence suggesting that protozoan parasites are able to manipulate host cells via epigenetic modifications of the host genome altering transcription and signaling pathways. This unexplored territory of epigenetic modulations in *C. parvum* infection would require further investigation for instance by ChIP-sequencing to identify all regulated genes.

## Materials and Methods

### *In silico* analysis

The protein sequences of the SET domains of several representative KMTs including *Saccharomyces cerevisiae* SET1 (GenBank Accession number EDN62358) and SET2 (NP012367), *Homo sapiens* EZH2 (NP004447), SUV39H1 (BAD96791), SET8 (NP065115), SMYD3 (NP001161212), *Toxoplasma gondii* KMTox (XP002371399) and AKMT (XP 002370918) were retrieved from databases and used as queries to search *C. parvum* homologs by performing BLASTp analysis on the database CryptoDB (http://Cryptodb.org). In parallel, JumonjiC (JmJC)-domain was also used for the search of lysine demethylases (KDMs) in the *C. parvum* genome. The presence of the conserved SET domain within the putative KMTs of *C. parvum* was confirmed by analyzing the identified sequences using the InterPro program (http://www.ebi.ac.uk) which integrates the signatures provided from 13 different databases (CATH, CDD, HAMAP, MobiDB Lite, PANTHER, Pfam, PIRSF, PRINTS, PROSITE, SFLD, SMART, SUPERFAMILY and TIGRFAMs). Furthermore, multiple sequence alignment was performed to compare the putative SET domain sequences of *C. parvum* with those of a panel of 31 representative KMTs using the MUltiple Sequence Comparison by Log-Expectation (MUSCLE) software under manual supervision. Simultaneously, a phylogenetic analysis was performed from the same set of SET domain sequences. All positions containing gaps and regions of ambiguous alignment were removed, yielding 116 sites for phylogenetic inference. Full-length alignment and boundaries can be available upon request to the corresponding author. Briefly, phylogenetic trees were constructed using the Neighbour-joining (NJ) and Maximum Likelihood (ML) methods implemented in Mega X (68) using the Jones-Taylor-Thornton (JTT) substitution model. The relative stability of topological elements was assessed using 1000 bootstrap replicates for both NJ and ML.

### Homology modelling

Three-dimensional (3D) models of SET domains identified in the *C. parvum* genome (CpSETs) were built with the automated comparative modeling program Swiss Model Interactive Workspace (https://swissmodel.expasy.org/interactive) using as homologous protein templates, highly resolved X-ray crystal structures of human SET1 (MLL1) (Protein Data Bank (PDB) code: 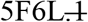; X-ray resolution 1.90 Å, SET2 (SETD2) (6J9J; 1.78 Å), and SET8 (SETD8) (5TEG; 1.30 Å). For each CpSET model developed, the quality of the structure was evaluated by the MolProbity web server (69). The MolProbity score is a combination of the clash score, rotamer and geometric parameters, and the Ramachandran evaluations into a single score (70). Lower MolProbity scores are better, meaning good quality structures. The MolProbity server reports also a percentile relative to the score distribution for crystal structures near the resolution of the submitted structure. In case of a modelled structure, the distribution is established covering all resolutions (range of 0Å-99Å). Distance matrix alignment (Dali) server (71) was used to perform pairwise structural alignment between the template and the newly generated CpSET models. The secondary structures were assigned using DSSP algorithm and the ChimeraX software was used to visualize the superimposition of templates and CpSET models (72).

### *Cryptosporidium* oocysts

Oocysts of *C*. *parvum* strain Iowa (purchased from Waterborne^™^, New Orleans, LA, USA) were stored in phosphate-buffered saline (PBS) with penicillin, streptomycin, gentamycin, amphotericin B and 0.001% Tween 20 at 4°C until use. Absence of bacteria and fungi was assured by testing the oocyst suspensions on both Plate Count Agar and Sabouraud plates at 37°C for 1 week. Oocysts viability was determined as previously described (73).

### *In vitro* culture

Human ileocecal adenocarcinoma cells (HCT-8; ATCC CCL-244) were maintained in Dulbecco’s modified eagle medium (DMEM) supplemented with 2 mM L-glutamine, 10% Fetal Bovine Serum (FBS) and antibiotics (100 U/ml penicillin, 100 μg/ml streptomycin). Cells were cultured at 37°C in humidified incubator supplemented with 5% CO2. Oocysts excystation was triggered as described previously (74). After infecting the cells with the parasite, the culture was maintained in RPMI 1640 medium supplemented with 2 mM L-glutamine, 15 mM HEPES buffer, 23 mM sodium bicarbonate, 5 mM glucose, 0.5 μM folic acid, 7 μM 4-aminobenzoic acid, 0.1 μM calcium pantothenate, 50 nM ascorbic acid, 1% (vol/vol) heat-inactivated fetal calf serum, 210 μM gentamycin, 170 μM streptomycin and penicillin (105 U/mL). For negative controls that received no parasites, only maintenance medium was applied onto the monolayers.

### Animal experiment

A total of 10 seven-week-old CB17-SCID mice were obtained from a colony bred at the Pasteur Institute of Lille (France). Mice were administered with 4 mg/L of dexamethasone (Merck, Lyon, France) through drinking water. Infective doses of *C. parvum* (10^5^ oocysts / mouse) were prepared as described previously (75) and inoculated by oral-gastric gavage. In order to quantify parasite shedding, mice faeces were collected and treated as described previously (65). At 60 days post-infection (PI) or when clinical signs of imminent death appeared, mice were euthanized by carbon dioxide inhalation. Experiments were conducted in the animal facility at the Institute Pasteur of Lille (research accreditation number, D 59 350 009). Animal protocols were approved by the French regional ethical committee with the number APAFIS#9621.

### Histopathology

Ileo-caecal regions were removed from each mouse, fixed in 4 % neutral formalin and embedded in paraffin. Sections of 4 μm thick were stained by hematoxylin-eosin-saffron (Leica Autostainer-XL, Rueil-Malmaison, France). Histological sections were analysed using a Leica DMRB microscope equipped with a Leica digital camera connected to an Imaging Research MCID analysis system (MCID software, Cambridge, United Kingdom). Neoplastic lesions at different sites were scored as previously described (75)

### Immunofluorescence assay

Sporozoites were fixed in 4% paraformaldehyde (PAF) for 10 min. After a wash with 1X PBS, the sporozoites were incubated in permeabilization solution (0.2 % Triton X-100 in 1X PBS) for 5 min then treated 10 min with blocking solution (0.3 M glycine, 1 % BSA, 0.1 % Tween 20 in 1X PBS). Finally, the sporozoites were incubated in primary antibody solutions for respective histone lysine methylations (Supplementary Table 1) overnight at 4 °C in a humidified chamber. The primary antibody solution was washed away with 1X PBS and the sporozoites were incubated with secondary antibody solution (Supplementary Table 1) for 1 h at room temperature. Following another wash with PBS, the sporozoites were incubated with the antibody *anti-Cryptosporidium* (Sporoglo, Waterborne^™^, New Orleans, LA, USA) for 45 min. After a final incubation with DAPI (1 μg/ml) for 15 min, the slides were mounted using Mowiol mounting medium (Mowiol ^®^ 4-88, Sigma, USA). For the *in vitro* staining, HCT-8 cells grown on coverslips in 24 well-plates were infected with 30,000 excysted oocysts per well and fixed at different time points PI: 6 h and 24 h to detect asexual stages and 55 h to detect sexual stages. The staining procedure was similar to that described above for sporozoites. For the *in vivo* staining, ileo-cecal sections of 5 μm thickness were obtained from formalin-fixed and paraffin-embedded specimens and placed on glass slides. The progressive rehydration was followed by an antigen retrieval step using citrate buffer pH 6.5 in a microwave oven for 15 min. After 1 h incubation in blocking buffer (2.5% BSA in 0.1 % Tween-20 1X PBS), the primary antibodies, diluted in blocking buffer, were applied for 1 h at 37°C. After three washes of 5 min with 1X PBS supplemented with 0.1 % Tween-20, the slides were incubated in the secondary antibodies for 1 h at 37°C. After a final wash, the slides were counterstained with DAPI and mounted with Mowiol mounting medium. The images were acquired using Zeiss LSM880 confocal microscope and analyzed using the ZEN lite Digital Imaging software.

### RNA extraction, cDNA synthesis and Real-Time quantitative PCR (RT-qPCR)

Total RNA was extracted from infected and non-infected HCT8 cells using NucleoSpin RNA Kit (Macherey-Nagel, Germany). An on-column DNase digestion with a RNase-free DNase was included in the process described by the fabricant to remove any genomic DNA contamination in RNA samples. RNA quality and quantity were determined using Agilent RNA6000 Nano kit by capillary electrophoresis (Agilent 2100 bioanalyzer, Agilent Technologies, Santa Clara, CA, USA). cDNA was synthesized from 1 μg of total RNA using oligo-dT primer and Superscript III reverse transcriptase (RT) in a 20 μl reaction (Invitrogen) according to the manufacturer’s instruction. Each amplification was performed in a volume of 20 μl containing 1 μl of cDNA, 200 nM of each primer and 1X Brilliant III Ultra-Fast SybrGreen qPCR Master Mix (Agilent Technologies). The RT-qPCR reactions were performed on a QIAGEN Rotor-Gene Q instrument (Corbett Research, Qiagen) and included an initial denaturation at 95°C for 3 min followed by a two-step cycling protocol consisting of 45 cycles of denaturation at 95°C during 10 s and annealing/extension at 60°C during 10 s. The PCR cycling program was followed by a standard melt step, stepwise increasing temperature each 5 s by 1°C, ranging from 65°C to 95°C. Primers used for RT-qPCR of putative CpKMTs are listed in supplementary Table 2. The 2^-ΔΔCt^ method was used to calculate the relative expression levels of *KMT g*enes with the constitutively expressed 18S rRNA gene as the internal reference and the ΔC_t_ value of the sporozoite stage as the calibrator.

### Purification of histones and Western blot

Parasite histones were enriched by performing a fractionation protocol. Briefly, HCT-8 cells infected and non-infected at different PI time points were incubated in ice cold fractionation buffer (25mM Tris-HCl pH 8.5, 50 mM NaCl, 0.1 % Triton-X100, 1 mM EDTA, 1x protease inhibitor (cOmplete^™^Protease Inhibitor Cocktail, Roche, USA). After dislocating the cells, the lysate was centrifuged at 2,000 g for 10 min at 4°C. Respective cellular fractions (pellet and supernatant) were subjected to histone purification using the EpiQuik^™^ Total Histones Extraction Kit (Epigentek, OP-0006-100, USA). The purified histone concentration was determined using micro BCA protein assay kit (Pierce, Thermofischer Scientific, USA). Approximately equal amounts of purified histones and parasite histones were separated by 15% sodium dodecyl sulfate polyacrylamide gel electrophoresis (SDS-PAGE) and transferred to nitrocellulose membranes (Millipore, USA). Chemiluminescent detection of bands was carried out by using Super Signal West Femto Maximum Sensitivity Substrate (Thermo Scientific, USA).

### Cloning and protein expression of CpSETs

To express the SET domains proteins, putative active domains of CpSET1 (aa residues 2100 – 2244) and CpSET8 (aa residues 402 – 556) were *de novo* synthetized and cloned into the bacterial expression vector pET15b at the NdeI and BamHI cloning sites which follows N-terminal 6x histidine tag sequence (Gencust, Boynes, France). The expression plasmid was amplified in *E. coli* BL21 (DE3) cells. After induction with 0.1 mM isopropyl β-D-1-thiogalactopyranoside (IPTG) at 16°C for 16 h, the cells were collected by centrifugation. The cell pellets were resuspended in lysis buffer (50 mM Tris-HCl, pH 7.5, 150 mM NaCl, 10 mM Imidazole, 1x protease inhibitor) then broken using bead beating method (Mixer Mill MM 400, Retsch, Germany). After centrifugation, his-tagged SET domains were recovered in the supernatant as soluble protein fraction.

### *In vitro* histone methyltransferase assay

To detect histone methyltransferase activity, 1 μg of recombinant H4 (New England Biolabs) and 100 μM S-adenosyl methionine (SAM) (Sigma, USA) were mixed with recombinant CpSET domains (CpSET8) in methyltransferase buffer (50 mM Tris-HCl (pH 8.5), 5 mM MgCl2, 4 mM DTT) at 30°C for 1 h. The reaction was stopped by adding 2x SDS sample buffer, and the reaction mixture was analyzed on 15% SDS-PAGE, followed by western blotting with antibodies against tri-methylation of lysine 20 of histone 4 (H4K20Me3) at a dilution of 1:1000 (Supplementary table 1).

### Signal quantification

Fluorescence intensity signals between infected and non-infected host tissue were quantified using ImageJ software version 1.52a (NIH, USA). Signal intensity was measured from individual nucleus of infected vs non-infected epithelial cell form intestinal crypts. For the statistical analysis, a mixed model was used to test the relationship between fluorescence intensity markers and group condition taking in account sample repetition. A mixed regression model was created considering fluorescence quantification as the main outcomes and sample identifier as random effect. The general significance level was set at a p-value below 0.05. All analyses were performed using packages nlme from the R statistical computing program (Version 4.1.1, date of release 8 October 2021; R Development Core Team, http://www.R---project.org, accessed on 12 January 2022) The data is represented using GraphPad Prism 9.1 (San Diego, California, USA)

## Acknowledgments

We wish to thank the Bioimaging Center Lille, Institut Pasteur de Lille, France for access to the microscopy instruments.

## Author contributions

Conceptualization: Manasi Sawant, Sadia Benamrouz-Vanneste, Jonathan Weitzman, Magali Chabé, Eric Viscogliosi, Gabriela Certad

Data curation: Manasi Sawant, Dionigia Meloni

Formal analysis: Manasi Sawant, Gaёl Even

Funding acquisition: Jonathan Weitzman, Gabriela Certad

Investigation: Manasi Sawant, Sadia Benamrouz-Vanneste, Dionigia Meloni, Nausicaa Gantois, Gaёl Even, Karine Guyot, Colette Creusy, Erika Duval, Magali Chabé

Methodology: Manasi Sawant, René Wintjens, Jonathan Weitzman, Magali Chabé, Eric Viscogliosi, Gabriela Certad

Project administration: Jonathan Weitzman, Eric Viscogliosi, Gabriela Certad,

Resources: Jonathan Weitzman, Eric Viscogliosi, Gabriela Certad,

Supervision:. Sadia Benamrouz-Vanneste, Jonathan Weitzman, Eric Viscogliosi, Gabriela Certad

Visualization: Manasi Sawant, Dionigia Meloni

Writing– original draft: Manasi Sawant, Sadia Benamrouz-Vanneste, Magali Chabé, Eric Viscogliosi, Gabriela Certad

Writing– review & editing: Manasi Sawant, Sadia Benamrouz-Vanneste, Dionigia Meloni, René Wintjens, Jonathan Weitzman, Magali Chabé, Eric Viscogliosi, Gabriela Certad

## Funding

This study was supported by the Plan Cancer “Epigénétique et cancer” 2015 (PARA-CAN #PARA-15-RCA), the Centre National de la Recherche Scientifique, the Institut National de la Santé et de la Recherche Médicale, the Institute Pasteur of Lille, the University of Lille and the Centre Hospitalier Regional Universitaire (CHRU) de Lille. M.S. was supported by a PhD fellowship from the University of Lille. R.W. is a Research Associate with the Belgian National Funds for Scientific Research (FRS-FNRS).

## Notes

### Competing Interest Statement

The authors have declared no competing interest.

