## Supplemental file for "Putative SET-domain methyltransferases in *Cryptosporidium parvum* and histone methylation during infection"

**Supplementary Figure 1:** Conserved sequences of histone proteins in Human, Mouse and *C. parvum*

```
             ....|....| ....|....| ....|....| ....|....| ....|....|
                      10         20         30         40         50
Hs H3.1      ARTKQTARKS TGGKAPRKQL ATKAARKSAP ATGGVKKPHR YRPGTVALRE
Hs H3.2      ARTKQTARKS TGGKAPRKQL ATKAARKSAP ATGGVKKPHR YRPGTVALRE
Hs H3.3      ARTKQTARKS TGGKAPRKQL ATKAARKSAP STGGVKKPHR YRPGTVALRE
Mm H3.1      ARTKQTARKS TGGKAPRKQL ATKAARKSAP ATGGVKKPHR YRPGTVALRE
Mm H3.2      ARTKQTARKS TGGKAPRKQL ATKAARKSAP ATGGVKKPHR YRPGTVALRE
Mm H3.3      ARTKQTARKS TGGKAPRKQL ATKAARKSAP STGGVKKPHR YRPGTVALRE
Cp H3        ARTKQTARKS TGGKAPRKQL ASKGARKSAP VTGGVKKPRR YRPGTVALRE


             ....|....| ....|....| ....|....| ....|....| ....|....|
                      60         70         80         90        100
Hs H3.1      IRRYQKSTEL LIRKLPFQRL VREIAQDFKT DLRFQSSAVM ALQEACEAYL
Hs H3.2      IRRYQKSTEL LIRKLPFQRL VREIAQDFKT DLRFQSSAVM ALQEASEAYL
Hs H3.3      IRRYQKSTEL LIRKLPFQRL VREIAQDFKT DLRFQSAAIG ALQEASEAYL
Mm H3.1      IRRYQKSTEL LIRKLPFQRL VREIAQDFKT DLRFQSSAVM ALQEACEAYL
Mm H3.2      IRRYQKSTEL LIRKLPFQRL VREIAQDFKT DLRFQSSAVM ALQEASEAYL
Mm H3.3      IRRYQKSTEL LIRKLPFQRL VREIAQDFKT DLRFQSAAIG ALQEASEAYL
Cp H3        IRRFQRSTEL LIRKLPFQRL VREIAQDFKT DLRFQSQAVM ALQEAAEAYL


             ....|....| ....|....| ....|....| ....|.
                     110        120        130
Hs H3.1      VGLFEDTNLC AIHAKRVTIM PKDIQLARRI RGERA.
Hs H3.2      VGLFEDTNLC AIHAKRVTIM PKDIQLARRI RGERA.
Hs H3.3      VGLFEDTNLC AIHAKRVTIM PKDIQLARRI RGERA.
Mm H3.1      VGLFEDTNLC AIHAKRVTIM PKDIQLARRI RGERA.
Mm H3.2      VGLFEDTNLC AIHAKRVTIM PKDIQLARRI RGERA.
Mm H3.3      VGLFEDTNLC AIHAKRVTIM PKDIQLARRI RGERA.
Cp H3        VGLFEDTNLC AIHAHRVTIM PKDIQLARRI RGER..


             ....|....| ....|....| ....|....| ....|....| ....|....|
                      10         20         30         40         50
Hs H4        SGRGKGGKGL GKGGAKRHRK VLRDNIQGIT KPAIRRLARR GGVKRISGLI
Mm H4        SGRGKGGKGL GKGGAKRHRK VLRDNIQGIT KPAIRRLARR GGVKRISGLI
Cp H4        SGRGKGGKGL GKGGAKRHRK VLRDNIQGIT KPAIRRLARR GGVKRISALI

             ....|....| ....|....| ....|....| ....|....| ....|....|
                      60         70         80         90        100
Hs H4        YEETRGVLKV FLENVIRDAV TYTEHAKRKT VTAMDVVYAL KRQGRTLYGF
Mm H4        YEETRGVLKV FLENVIRDAV TYTEHAKRKT VTAMDVVYAL KRQGRTLYGF
Cp H4        YEEVRGVLKA FLETVIKDAV TYTEYARRKT VTAMDVVHAL KRQGKTLYGF

             ....|....|
                     110
Hs H4        GG........
Mm H4        GG........
Cp H4        GG........
```

**Supplementary Figure 1:** Multiple sequence alignment of histone proteins from homo sapiens, Mus musculus and *C. parvum.* Highly conserved lysine residues (K) are highlighted in red. The uniport accession numbers of the sequences used are Hs H3.1 (P68431), Hs H3.2 (Q71DI3), Hs H3.3 (P84243), Hs H4 (Q5CV68), Ms H3.1 (P84228), Ms H3.2 (P84244), Ms H3.3 (P68433), Ms H4 (P62806), Cp H3 (Q5CUJ9) and Cp H4 (Q5CV68)

**Supplementary Figure 2** : Western blot analysis of parasite histone methylation marks.

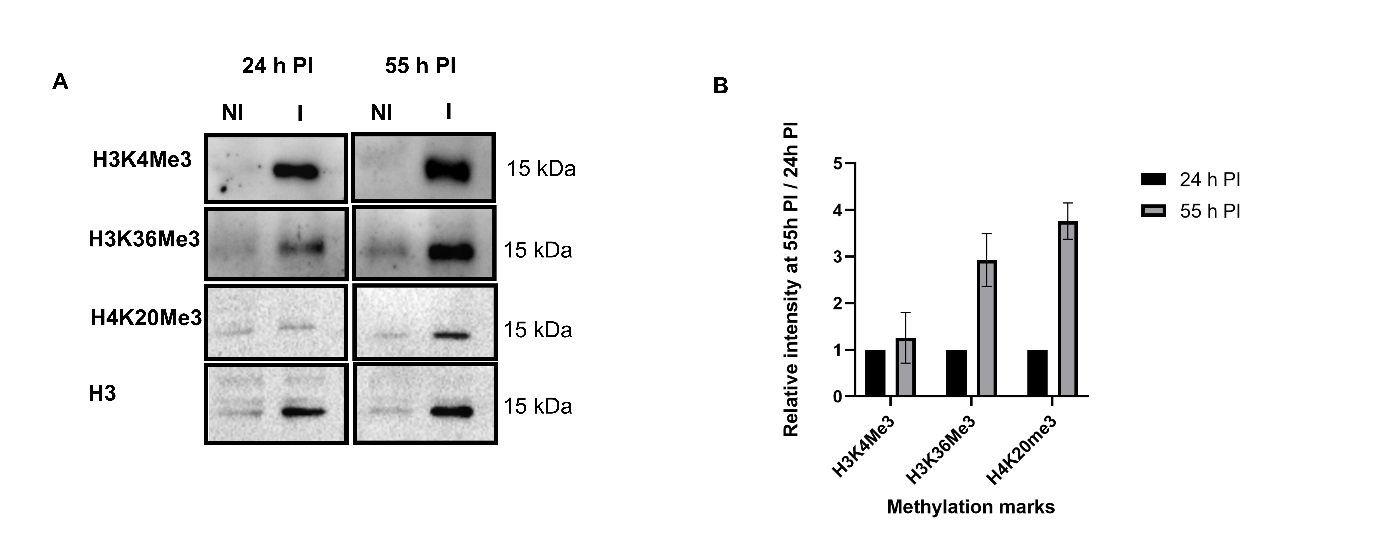

A. Western blotting analysis of histone lysine methylation modifications during *C. parvum* development *in vitro* at 24 h (asexual stages) and 55 h (sexual stages) PI after purification of histones from the parasites. B. The histograms represent the relative intensity signals of methylation marks in the parasite at 55h PI relative to 24h PI. Each sample was normalized to the H3 used as internal control. The graph represents means in triplicate values. NI – Non-infected HCT-8 cells. I – Infected HCT-8 cells. Scale bar – 1 µm. Results are representative of three independent experiments.

**Supplementary Figure 3** : Quantification of fluorescence intensity signals of anti-methylation antibodies in *C. parvum* infected vs non-infected ileo-caecal tissue.

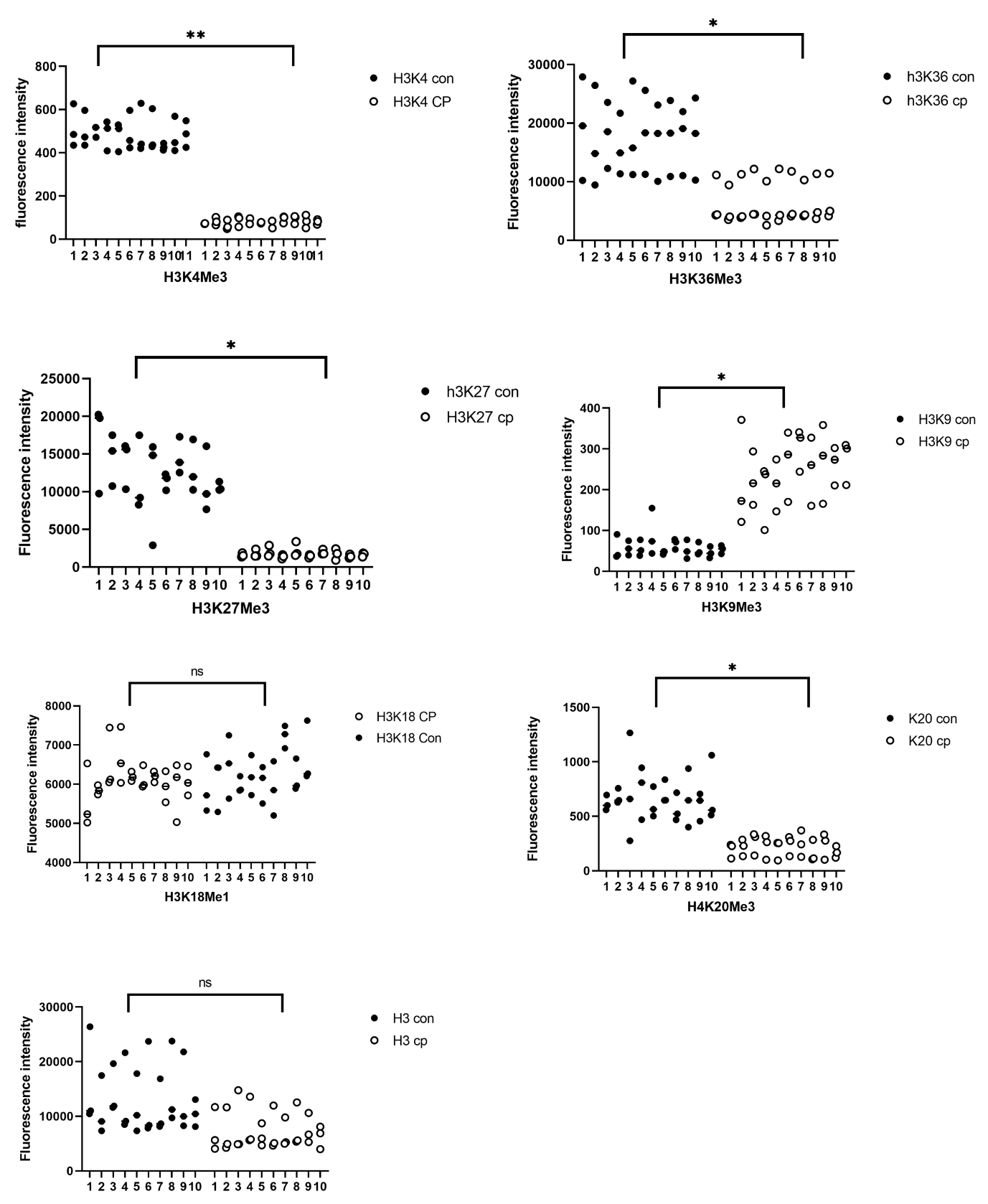

**Supplementary Figure 3** – Fluorescence intensity signals of anti-H3K4Me3, H3K36Me3, H3K27Me3, H3K9Me3, H3K18Me1, H4K20Me3 antibodies in *C. parvum* infected vs non-infected ileo-caecal tissue were quantified. The signal intensities were measured by nuclei. Numbers on the Y axis indicate individual nucleus. The black circles indicate uninfected nuclei. The white circles indicate infected nuclei. For the statistical analysis, a mixed regression model was created considering fluorescence quantification  as the main outcomes and sample identifier as random effect. Significance was determined for p<0.05. Con – Non infected tissue, Cp – Infected tissue. The experiments were repeated at least three times.

**Supplementary Figure 3**: Relative intensity signal in infected HCT-8 cells vs non-infected

HCT-8 cells.

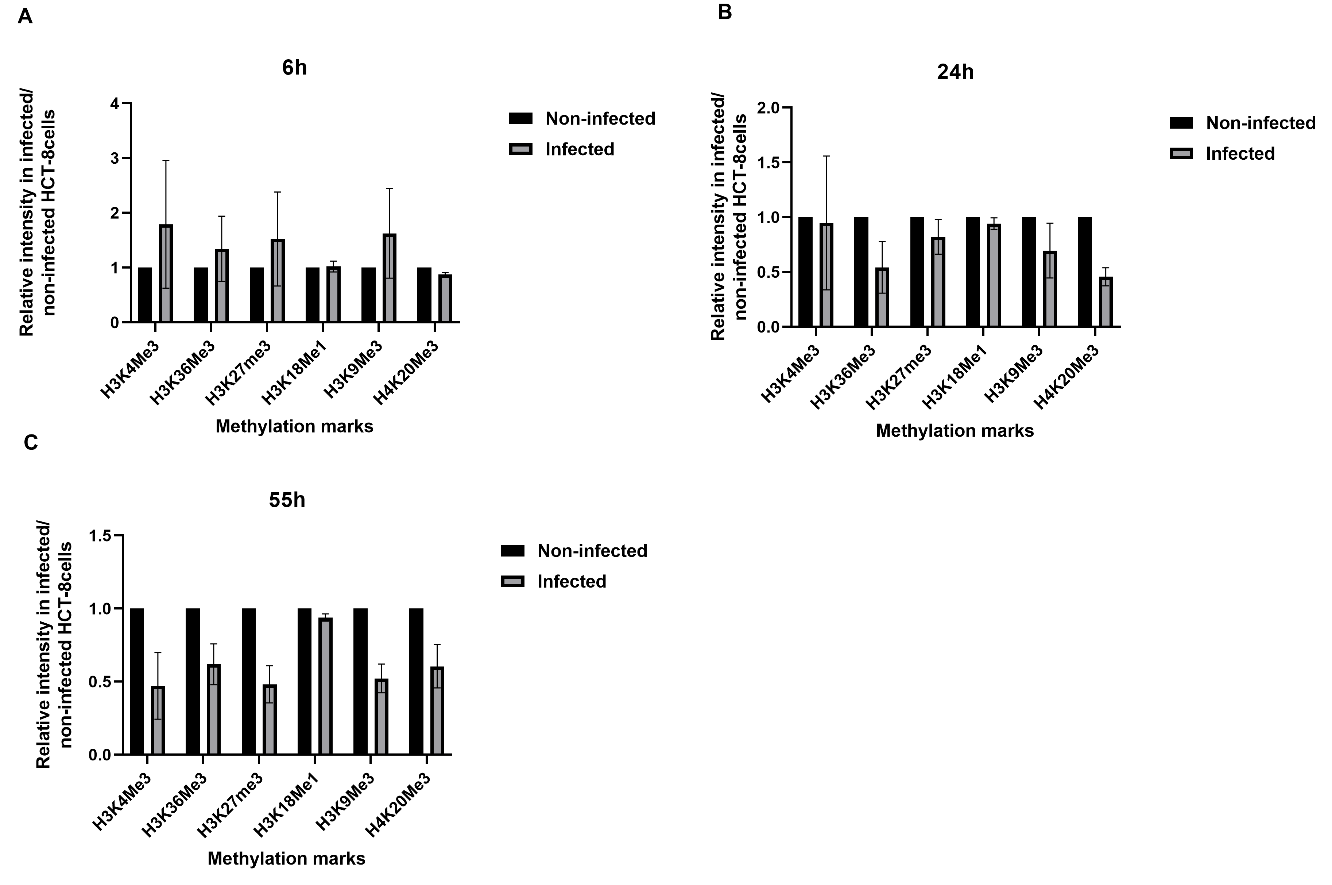

**Supplementary Figure 3** - The histograms represent the relative intensity signals of methylations in the host cells. The relative intensity signal was measured in infected HCT-8 cells with respect to non-infected HCT-8 cells. Each sample was normalized to the H3 used as internal control. The graph represents means in triplicate values.

**Supplementary Table 1**: Primary and secondary antibodies dilutions

| Antibodies | ICC/ IHC (dilutions) | Western blotting (dilutions) |
| --- | --- | --- |
| Anti-Histone H3 (tri methyl K4) antibody - ChIP Grade ab8580 | 1:1000 | 1:1000 |
| Anti-Histone H3 (tri methyl K36) antibody - ChIP Grade ab9050 | 1:500 | 1:1000 |
| Anti-Histone H3 (tri methyl K9) antibody - ChIP Grade (ab8898) | 1:500 | 1:1000 |
| Anti-Histone H3 (tri methyl K27) antibody - ChIP Grade [mAbcam 6002] | 1:500 | 1:1000 |
| Anti-Histone H3 (mono methyl K18) antibody [EPR17710] | 1:5000 | 1:20000 |
| Anti-Histone H4 (tri methyl K20) antibody [EPR17001(2)] - ChIP Grade | 1:1000 | 1:1000 |
| Anti-Histone H3 antibody - Nuclear Marker and ChIP Grade [ab1791] | 1:100 | 1:1000 |
| Goat Anti-Rabbit IgG H&L (Alexa Fluor® 488)(ab150077) | 1:1000 |  |
| A600FLR-20X Sporo-Glo™ | 1:50 |  |
| Goat Anti-Rabbit IgG H&L (HRP) (ab6721) |  | 1:1000 |

Abbreviations: IHC: immunohistochemistry; ICC: Immunocytochemistry

**Supplementary Table 2** : Primers used for RT-qPCR analysis of putative KMTs

| **Gene ID** | **Primers** | **Tm** | **Sequences** | **Fragment**  **size (bp)** |
| --- | --- | --- | --- | --- |
| cgd1-2170 | F | 60.1 | 5’ gctgaagcagtatcccgttgca 3’ | 80 |
|  | R | 58.2 | 5’ tcgtctttcatacccagttcttgc 3’ |  |
| cgd4-370 | F | 58.4 | 5’ tgtgtaatcgccggatctc 3’ | 91 |
|  | R | 59.7 | 5’ gctctttggcctcgttaagc 3’ |  |
| cgd4-2090 | F | 59.9 | 5’ gccagggaatttgggtttaacg 3’ | 84 |
|  | R | 59.7 | 5’ ttcatcggttgcaatccctcc 3’ |  |
| cgd5-400 | F | 58.9 | 5’ gaaagatcctgcggagtatgc 3’ | 116 |
|  | R | 58.5 | 5’ tcttcgagtccgacgca 3’ |  |
| cgd5-2340 | F | 58.2 | 5’ tgctaacgatggaagcgca 3’ | 84 |
|  | R | 58.3 | 5’ gatctaccttctctttcgtcataccac 3’ |  |
| cgd6-1470 | F | 58.3 | 5’ gtagcttgcctagattggaaagca 3’ | 92 |
|  | R | 59.5 | 5’ tggaataagtcctgttcctagctc 3’ |  |
| cdg7-5090 | F | 58.6 | 5’ agtgaatccacgacaaaaagctcc 3’ | 80 |
|  | R | 58.4 | 5’ cccatccaagaatgcttgg 3’ |  |
| cgd8-2730 | F | 58.2 | 5’ tgacggtagaaagtgctagga 3’ | 116 |
|  | R | 58.8 | 5’ cttgcttgatgaggaatgagagc 3’ |  |
| 18S | F | 58.2 | 5’ tgccttgaatactccagcatgg 3’ | 103 |
|  | R | 59.6 | 5’ tacaaatgcccccaactgtcc 3’ |  |
